# Crosstalk between tubulin glutamylation and tyrosination regulates kinesin-3-mediated axonal transport

**DOI:** 10.1101/2025.08.09.669510

**Authors:** Odvogmed Bayansan, Muhammad Safwan Khawaja, Dwika Sarnia Putri, Oliver Ingvar Wagner

## Abstract

Tubulin post-translational modifications (PTM) are critical regulators of microtubule function and diversity in neurons. In *Caenorhabditis elegans*, we investigated how tubulin deglutamylases (CCPP-1 and CCPP-6) and polyglutamylases (TTLL-5 and TTLL-9) affect kinesin-3 KIF1A/UNC-104–mediated axonal transport. Loss of CCPP-1 not only facilitates tubulin polyglutamylation as expected, but it also leads to increased tyrosination in Western blots. Vice versa, loss of TTLL-5 reduces tubulin polyglutamylation as expected, but also leads to reduced tyrosination signals. This crosstalk in tubulin PTM appears to be a critical feature as no tyrosination or detyrosination enzymes are known in *C. elegans*. Notably, acetylation and detyrosination remain unaffected in the deglutamylation and polyglutamylation mutants. Functionally, reduced glutamylation and tyrosination improved UNC-104 motility. Conversely, increased glutamylation and tyrosination negatively affected the movement of both the UNC-104 motor and its cargo RAB-3. UNC-104 motors visibly accumulate in neuronal cell bodies of *ccpp-1* mutants while being significantly reduced in *ttll-5* mutants. In *ttll-5* mutants, motors tend to cluster along distal axonal regions and these clusters are reduced in *ccpp-1* mutants revealing a role of tubulin PTM in axonal motor scaffolding. Employing promoter fusions, we confirmed that all investigated PTM enzymes express in neurons and colocalize with UNC-104. Moreover, co-immunoprecipitation assays revealed that hyperglutamylated tubulin appears in a physical complex with UNC-104, while hypoglutamylated tubulin binds less effectively to the motor. In our model, highly negatively charged polyglutamylated tubulin traps UNC-104 onto microtubules via increased charge-interactions. Tubulin “stickiness” is reduced in polyglutamylase mutants leading to increased motor speeds. Reduced synaptic vesicle transport in *ccpp-1* mutants has a negative impact on the nematode’s touch sensing, highlighting *C. elegans* as a valuable model for investigating tubulin PTM-related neurological disorders.

## INTRODUCTION

Axonal transport is a fundamental cellular process essential for neuronal function and survival. Two major classes of microtubule-based molecular motors drive this transport: kinesins mediate anterograde transport (from cell bodies to distal synapses), while cytoplasmic dynein is responsible for retrograde transport (from synapses back to the cell body). Axonal transport requires precise regulation through multiple mechanisms, including motor protein activation/inactivation, cargo binding specificity, and coordination between opposing motors, known as tug-of-war. Disruptions in axonal transport are linked to numerous neurodegenerative disorders, including Alzheimer’s disease, Parkinson’s disease, amyotrophic lateral sclerosis (ALS), as well as Huntington’s disease (Millecamps and Julien, 2013). Kinesin-3 KIF1A is the major axonal transporter of synaptic vesicles and is termed UNC-104 in *C. elegans*. KIF1A/UNC-104 comprises a N-terminal motor domain, a regulatory neck linker, a lysine-rich K-loop, a forkhead-associated (FHA) domain, coiled-coil regions that mediate dimerization, and a C-terminal cargo-binding domain containing a pleckstrin homology (PH) domain for membrane attachment. The K-loop, a positively charged region enriched in lysine, serves as a microtubule-binding surface by interacting with the glutamate-rich and negatively charged C-terminal side chains of β-tubulin (E-Hook) (Lessard et al., 2019; Soppina and Verhey, 2014). Mutations in the unc-104 gene result in severe paralysis of nematodes due to synaptic transmission defects (Hall and Hedgecock, 1991), and KIF1A-deficient mice die during embryonic stages due to sensory neuron degeneration (Yonekawa et al., 1998). Mutations in KIF1A are linked to hereditary spastic paraparesis (HSP), Charcot-Marie-Tooth (CMT) disease and KIF1A-associated neurological diseases (KAND) (Citterio et al., 2015; Morikawa et al., 2022). UNC-104 motor activity and cargo recognition are tightly regulated by synaptic scaffolding proteins Liprin-α/SYD-2, CASK/LIN-2 (Muniesh et al., 2020; Wagner et al., 2009; Wu et al., 2016) and RIM/UNC-10 (Bayansan et al., 2025). Although microtubules are structurally conserved, they exhibit remarkable functional diversity across different cell types due to the incorporation of various tubulin isotypes as well as rich post-translational modifications (PTM) (Janke and Magiera, 2020). Tubulin PTMs regulate the interaction of microtubules with microtubule-associated proteins (MAP), motor proteins, and microtubule severing enzymes (e.g., spastin), all of them modulating microtubule dynamics, intracellular transport and cell polarity. Mutations in tubulin isotypes result in disruption of the microtubule network during early brain development, leading to a group of neurological disorders known as tubulinopathies. Tubulin PTMs are mediated by enzymes, occurring either inside the tubulin body (e.g., acetylation) or on the flexible C-terminal tails (e.g., glutamylation or tyrosination) (Janke and Magiera, 2020). Tubulin polyglutamylation involves the addition of negatively charged glutamate chains to the C-terminal tails of α– and β-tubulin by tubulin tyrosine ligase-like (TTLL) enzymes. The mammalian genome encodes nine TTLL enzymes (TTLL1, 2, 4, 5, 6, 7, 9, 11 and 13) (van Dijk et al., 2007) with TTLL1 and 7 abundantly expressed in neurons. TTLL1 preferably modifies α-tubulin, while TTLL7 modifies β-tubulin. TTLL1 acts to prevent degeneration of Purkinje cells (*pcd*) and myelinated axons in mice (Bodakuntla et al., 2021). Polyglutamylation regulates binding between kinesins, MAP1B, MAP2 and Tau (Bonnet et al., 2001), and loss of TTLL1 leads to decreased microtubule-binding affinity of KIF1A, but not KIF3A or KIF5 (Ikegami et al., 2007). Increased polyglutamylation is also correlated to increased microtubule severing by recruiting spastin and katanin (Lacroix et al., 2010; Valenstein and Roll-Mecak, 2016). The *C. elegans* genome encodes six TTLLs: TTLL-4, –5, –9, –11,-12, and –15 (Lu and Zheng, 2022). TTLL-4 is homolog to TTLL4, TTLL-5 is homolog to TTLL5, TTLL-9 is homolog to TTLL9, TTLL-11 is homolog to TTLL11, and TTLL-15 is ortholog to TTLL5 (Chawla et al., 2016). Polyglutamylation chains are removed by *c*ytosolic *c*arboxy*p*eptidase enzymes (CCP) (Kalinina et al., 2007; Rogowski et al., 2010). Six CCP enzymes have been identified in mammals (Berezniuk et al., 2013; Wu et al., 2015); however, only CCP1, 4, 5 and 6 reveal functional deglutamylase activity (Kimura et al., 2010). CCP1 removes penultimate glutamate residues from α-tubulin to produce Δ2 tubulin (Berezniuk et al., 2012; Hotta et al., 2023). While CCP1 shortens polyglutamylation chains from the C-terminus of tubulin by removing long glutamylation chains from α– and β-tubulin, CCP5 preferentially cleaves off α– and γ-linked glutamates at the branch point of tubulin (Berezniuk et al., 2013; Rogowski et al., 2010). Absence of CCP1 leads to hyperglutamylated tubulin, resulting in neuronal degeneration, axonal swelling and perturbed axonal transport (Fernandez-Gonzalez et al., 2002; Magiera et al., 2018). *C. elegans* expresses only two deglutamylase enzymes, CCPP-1 and 6 (Kimura et al., 2010; O’Hagan et al., 2011). CCPP-1 is homolog to CCP1, while CCPP-6 shares homology with CCP5 in mammals (Kimura et al., 2010; O’Hagan et al., 2011). In male specific neurons, loss of CCPP-1 causes defects in ciliary structure and regulates localization and motility of kinesin-3 KLP-6 and PKD-2 as well as kinesin-2 KIF17/OSM-3. In injured *C. elegans* neurons, DLK-1 signaling controls microtubule dynamics by downregulating KLP-7 (kinesin-13) and upregulating CCPP-6. The latter enhances Δ2 modification of α-tubulin, thereby promoting microtubule growth and sustaining axon regeneration (Ghosh-Roy et al., 2012). Tubulin tyrosination is catalyzed by tubulin tyrosine ligase (TTL) enzymes that add tyrosination residues to the C-terminus of detyrosinated α-tubulin. Tyrosine can be removed by metalloprotease vasohibin (VASH1/2) in complex with small vasohibin-binding protein (SVBP) (Aillaud et al., 2017). Detyrosinated tubulin can then undergo further processing involving the removal of the final two or three C-terminal glutamate residues from α-tubulin via CCP generating (neuron specific) Δ2 and Δ3 tubulins. Tyrosinated tubulin is a marker of dynamic microtubules in growth cones and distal regions of neurites. In contrast, detyrosinated tubulin is a feature of stable microtubules in axons (Moutin et al., 2021). Kinesin-1 motor preferably binds to detyrosinated microtubules in axons (Dunn et al., 2008), while kinesin-3 and kinesin-5 preferably bind to tyrosinated tubulin in regions with dynamic microtubules present (growth cones, cell body) (Guardia et al., 2016; Kahn et al., 2015). Absence of tubulin tyrosination enzymes induces lethal neuronal defects in mice (Pagnamenta et al., 2019). Interestingly, neither tyrosination nor detyrosination enzymes are expressed in *C. elegans*. Acetylation occurs at lysine 40 of α-tubulin inside the hollow microtubule body and is catalyzed by the enzyme α-tubulin acetyltransferase (αTAT) homologous to MEC-17 in *C. elegans*. This modification is prevalent in long-lived microtubules to enhance microtubule stability and resistance to mechanical stress (Li and Yang, 2015). Loss of MEC-17 leads to microtubule instability and axonal degeneration in *C. elegans* (Neumann and Hilliard, 2014).

## MATERIALS AND METHODS

### Worm strains and plasmid design

Worms are maintained at 20°C on NGM agar plates seeded with an uranyl auxotroph OP50 *E. coli* strain according to standard methods (Brenner, 1974). Rescue strain *e1265*;*Punc-104::unc-104::mrfp* and construct design of *Punc-104::unc-104::mrfp* as well as *Punc-104::mCherry::rab-3* were described previously (Tien et al., 2011; Wagner et al., 2009). To generate polyglutamylase mutants expressing UNC-104::mRFP, we crossed *e1265*;*Punc-104::unc-104::mRFP* male worms with either OC343 *ttll-5(tm3360)* or OC419 *ttll-9(tm3889)* hermaphrodites to generate OIW 128 *ttll-5(tm3360);nthEx[Punc-104::unc-104::mrfp]* and OIW 129 *ttll-9(tm3889);nthEx[Punc-104::unc-104::mrfp],* respectively. To generate deglutamylase mutants expressing UNC-104::mRFP, we crossed *e1265*;*Punc-104::unc-104::mrfp* male worms with either RB1522 *ccpp-1(ok1821)* or RB625 *ccpp-6(ok382)* hermaphrodites to generate OIW 130 *ccpp-1(ok1821);nthEx[Punc-104::unc-104::mrfp]* and OIW 131 *ccpp-6(ok382);nthEx[Punc-104::unc-104::mrfp]*, respectively. Double mutants were generated by crossing *ccpp-6(ok382);nthEx[Punc-104::unc-104::mrfp]* male worms with *ccpp-1(ok1821)* hermaphrodites to generate OIW 131 *ccpp-1(ok1821);ccpp-6(ok382);nthEx[Punc-104::unc-104::mrfp]* animals. For all crossings, homozygous animals were screened after backcrossing and then genotyped. For cargo motility experiment, existing plasmid *Punc-104::mCherry::rab-3* was injected at 80 ng/µl into N2, *ccpp-1(ok1821)* and *ccpp-6(ok382)* mutants to either generate OIW 132 N2;*nthEx[Punc-104::mcherry::rab-3],* OIW 133 *ccpp-1(ok1821);nthEx[Punc-104::mcherry::rab-3]* or OIW 134 *ccpp-6(ok382);nthEx[Punc-104::mcherry::rab-3]*. For transcriptional fusions, we amplified the promoter region of ccpp-1 using forward aaaGCATGCgaagctgaaaagttg and reverse CTTCTTTACAGTATTTCAGCTGaaa primers to generate a 2534 bp amplicon. ccpp-6 region was amplified using forward aaaaGCATGCtgaacgtgaaataaatc and gttttaaagtaaaagttaCAGCTGaaa reverse primers to generate a 3512 bp amplicon. The transcriptional region of ttll-5 was amplified using forward aaaAAGCTTtttttcatcacactcgg and GAGGATTTTTGACGACGTCaaa reverse primers to generate a 2588 bp amplicon. pPD95.77 vector from Andrew Fire library was then used to generate *Pccpp-1::gfp, Pccpp-6::gfp* and *Pttll-5::gfp*. For colocalization assays, we injected either plasmid *Pccpp-1::gfp*, *Pccpp-6::gfp* or *Pttll-5::gfp* along with *Punc-104::unc-104::mrfp* (at each 80 ng/µl) into N2 wild type worms to generate OIW 134 N2;*nthEx[Punc-104::unc-104::mrfp;Pccpp-1::gfp],* OIW 135 N2;*nthEx[Punc-104::unc-104::mrfp;Pccpp-6::gfp]* or OIW 136 N2;*nthEx[Punc-104::unc-104::mrfp; Pttll-5::gfp]*.

#### Western blotting and Co-immunoprecipitation assays

Young adult worms from 15-20 medium-sized plates were used for protein extraction, following a published protocol (Ward et al., 1988). Worms were harvested by washing the plates thrice with 0.1 M NaCl solution, followed by 60% sucrose flotation. Worms from the sucrose layer were collected and washed twice with cold water and 1x homogenization buffer (HB). Subsequently, 1x HB with DTT and protease inhibitor was added to the harvested worms and the sample was stored at –80°C overnight. The following day, harvested worms were subjected to French press for protein extraction. Quantification was then carried out using commercial BCA kits (Prod# 23225, Pierce BCA protein kit, Thermo Scientific). For Western blotting, 100 µg of total protein was loaded per lane. Membranes were blocked with milk powder for 1 hour at room temperature. To detect polyglutamylated tubulin (PolyE), we used anti-Polyglutamylate chain (AG-25B-0030, AdipoGen, polyclonal, rabbit) antibody at 1:2000 dilutions, followed by anti-rabbit secondary antibody (GTX 213110-01) incubation at 1:4000 dilution. Tyrosinated tubulin was detected using an anti-tyrosinated tubulin antibody (T9028, Sigma, monoclonal, mouse) at 1:1000 dilutions, followed by an anti-mouse secondary antibody (GTX 213111-01) at 1: 2000 dilution. Detyrosinated tubulin was detected using anti-detyrosinated tubulin (AB3201, Sigma, polyclonal, rabbit) at 1:500 dilutions, followed by anti-rabbit secondary antibody (GTX 213110-01) at 1:1000 dilutions. Acetylated tubulin was detected using anti-acetylated tubulin T7451 (6-11B-1, Thermo Fisher, monoclonal, rabbit) at 1: 500 dilutions, followed by anti-rabbit secondary antibody (GTX 213110-01) incubation at 1:1000 dilutions. Anti-GAPDH antibody (GTX100118, GeneTex, polyclonal, rabbit) at 1:2500 dilutions, followed by anti-rabbit secondary antibody (GTX 213110-01) incubation at 1:5000 dilution was used as a positive control. Primary antibodies were incubated at 4°C overnight while secondary antibodies were incubated at room temperature for 4 hours. To detect lower molecular weight proteins, we used a 12% SDS-PAGE gel with 30% acrylamide, while for higher molecular weight proteins, a 6% SDS-PAGE gel with 40% acrylamide was used. For coimmunoprecipitation (Co-IP) assays, 1 mg protein extract was incubated with 50 µl protein G beads (CAT# LSKMAGG10, Millipore) and 5 µg UNC-104 antibody (GeneTex, commissioned by us, detects a N-terminal epitope, polyclonal, rabbit). After overnight incubation, worm lysates were subjected to Western blotting using polyglutamylated antibody (PolyE) to detect polyglutamylated tubulin bound to UNC-104. Antibody dilutions and incubation times were identical to those described for Western blot assays. All band intensity measurements were performed using NIH ImageJ software, following a published protocol (Schneider et al., 2012).

### Motility and motor clustering analysis

Worms were immobilized on 2% agarose-coated cover slides along with 7 µl of M9 buffer for motility and motor clustering analysis experiments. No anesthetics (such as Levamisole) were used. An Olympus IX81 microscope with a DSU Nipkow spinning disk unit connected to an Andor iXon DV897 EMCCD camera was employed for high-resolution and high-speed time-lapse imaging at 4-5 frames per second. Time-lapse images from a selected area were captured and then converted to kymographs using imaging analysis software NIH ImageJ. The “straighten plugin” was used to straighten curved axons and the “reslice” and “stack” plugins were used to generate kymographs (Suppl. Fig. S1B). A pause is defined if motors or vesicles move less than 0.07 µm/s, and each calculated velocity event does not contain any pauses. A moving event is defined as a single motility occurrence typically right after a pause or a reversal, and such an event ends when the motor again pauses or reverses (Suppl. Fig. S1C). For motor clustering analysis, ALM neurons were divided into four regions: cell body, initial, middle and distal regions as shown in Suppl. Fig. S1D. The number of particles in cell bodies was counted manually by visual inspection. For axonal regions, the NIH ImageJ “analyze particles” plugin was used to quantify the number of clusters in axons. After straightening the image, the threshold was adjusted and following parameters applied: distance in pixels = 1, known distance = 0.159, pixel aspect ratio = 1, and unit of length = µm. Travel distances of UNC-104 clusters from axon hillock to initial region were measured using the NIH ImageJ “line tool” after straightening the neuron and applying the same parameters (Figs. 7C, 8F). Density of UNC-104 clusters was calculated by dividing the number of particles in different regions of the axons by 10 μm (Figs. 7B, 8E). PMUV (persistent motility at uniform velocities) refers to how long a motor continues to move at a constant speed (Figs. 6G, 8C). For correlation analysis in Fig. 7D-G, r-values were calculated within two selected datasets (x = velocity, y = distance) using Pearson’s correlation coefficient and unpaired, two-tailed Student’s t-test to determine the p values.

### Confocal imaging

Worms were immobilized on 2% agarose pads with 7 µl of M9 and covered with a cover slide for imaging. No anesthetics (such as Levamisole) were used. A Zeiss LSM780 confocal laser scanning microscope was employed for fluorescent imaging (Fig. 9, Suppl. Fig. S2). To quantify colocalization between UNC-104 and promoter fusions of genes encoding PTM enzymes, we employed both Mander’s (MOC) and Pearson’s (PCC) correlation coefficients. MOC ranges between 0 and 1 and PCC ranges between –1 and 1. MOC measures the co-occurrence or overlap of two signals, based on absolute pixel intensities. PCC determines the linear correlation between pixel intensities, independent of absolute intensity values. The NIH ImageJ “polygonal selection tool” was selected to determine the region of interest (ROI) in an image. After background subtraction, NIH ImageJ “intensity correlation analysis” plugin was used to generate either MOC or PCC. PDM images refer to the product of the differences from the mean of red and green fluorescence intensities. It is calculated as: (red intensity – mean red intensity) × (green intensity – mean green intensity).

#### Whole-mount immunostaining

Young adult *e1265*;Punc-104::UNC-104::GFP rescue worms were synchronized, washed with M9 buffer, and collected in 2 ml Eppendorf tubes. Worms were then fixed with 1 ml of 4% paraformaldehyde and subjected to three cycles of freeze-thawing in liquid nitrogen, each lasting 3 minutes. After the final freeze cycle, worms were incubated in 4% paraformaldehyde at 4°C overnight. The following day, worms were washed with PBST and incubated with 1 ml of β-mercaptoethanol at 37°C overnight. After incubation, worms were once more washed with PBST and treated with 500 µl of collagenase (Cat# C0773, Sigma) at 37°C for 6.5 hours. Worms were then blocked for 1 hour at room temperature. Afterwards, worms were washed with antibody buffer and incubated with primary antibodies (either anti-Polyglutamylated tubulin or anti-tyrosinated tubulin) at 1:200 dilutions (4°C overnight). Subsequently, worms were incubated with secondary antibodies (anti-rabbit ab150119 or anti-mouse A-11005) at 1:500 dilutions (4 hours at room temperature). Worms were then washed thrice with antibody buffer, with each washing lasting 15 min. Afterwards, worms were mounted on a microscope slide and subjected to imaging.

#### Harsh and soft touch responses

To investigate the physiological consequences of the mutations, we use N2 and mutant worms at young adult stages. Both soft and harsh touch assays were performed to assess touch responses. For the soft touch assay, an eyelash was used to gently strike the worm laterally at the mid-body region. In contrast, a platinum wire was used for the harsh touch assay, applying a stronger stimulus to the same area. Each worm was tested four times per assay. A positive response was defined as a clear backward movement following the stimulus (MEC response). Worms that responded as expected all four times were scored as 100%, indicating full responsiveness (Chalfie, 2014).

#### Statistics

To determine whether acquired data follow a Gaussian distribution or not, we plotted histograms. If a Gaussian distribution was present, we used ordinary one-way ANOVA with Dunnett’s test for multiple comparisons or unpaired Student’s t-test for two-group comparisons. For non-Gaussian distributions, we used either (non-parametric) Kruskal-Wallis one-way ANOVA Dunn’s test for multiple comparisons, or unpaired Mann-Whitney test for two-group comparisons. For Western blot and Co-IP data analyses, we used either one-way ANOVA with Dunnett’s test for multiple comparisons, or unpaired two-tailed Student’s t-test for two-group comparisons. Box-and-whisker plots display the maximum, upper quartile, median, lower quartile, and minimum values.

## RESULTS

### Distinct effects of CCPP-1 and CCPP-6 on tubulin glutamylation

To investigate the effects of ccpp-1 and ccpp-6 mutations on tubulin glutamylation, we employed anti-polyglutamylated tubulin PolyE antibody, which detects polyglutamylation chains of at least four glutamate residues on tubulin C-terminal tails. In mammals, deglutamylase CCP1 effectively shortens glutamylated C-terminal chains from α– and β-tubulin (Berezniuk et al., 2012). To examine whether a similar function exists for CCPP-1 in *C. elegans*, we conducted Western blots, using strain RB1522 *ccpp-1(ok1821)* that carries a 1770 bp deletion in putative 5′ regulatory sequences and the first 5 exons of the *ccpp-1* gene, resulting in a null mutant (O’Hagan et al., 2011) (Fig. 1A). Indeed, the level of polyglutamylated tubulin in animals carrying the *ok1821* allele was significantly elevated (Fig. 1B-D). In *C. elegans*, CCPP-6 is the homolog of mammalian CCP5 and unlike CCP1, it specifically removes the glutamate at the branching-point; meaning, the first glutamate directly linked to the γ-carboxyl group of the glutamate residue in the tubulin backbone (Berezniuk et al., 2013). To investigate the function of CCPP-6, we use strain RB625 *ccpp-6(ok382)* that carries a 532 bp deletion removing the start codon of the gene as well as the first two exons (Dominguez et al., 2022) (Fig. 1A). From Figure 1B it is obvious that the level of polyglutamylated tubulin remains unaffected in in *ccpp-6(ok382)* mutants (Fig. 1 B+C and E). Interestingly, in *ccpp-1/ccpp-6* double mutants hyperglutamylation occurs (Fig. 1 B+C and F) pointing to synthetic interactions of these genes in parallel (and partially redundant) pathways.

**Figure 1:**
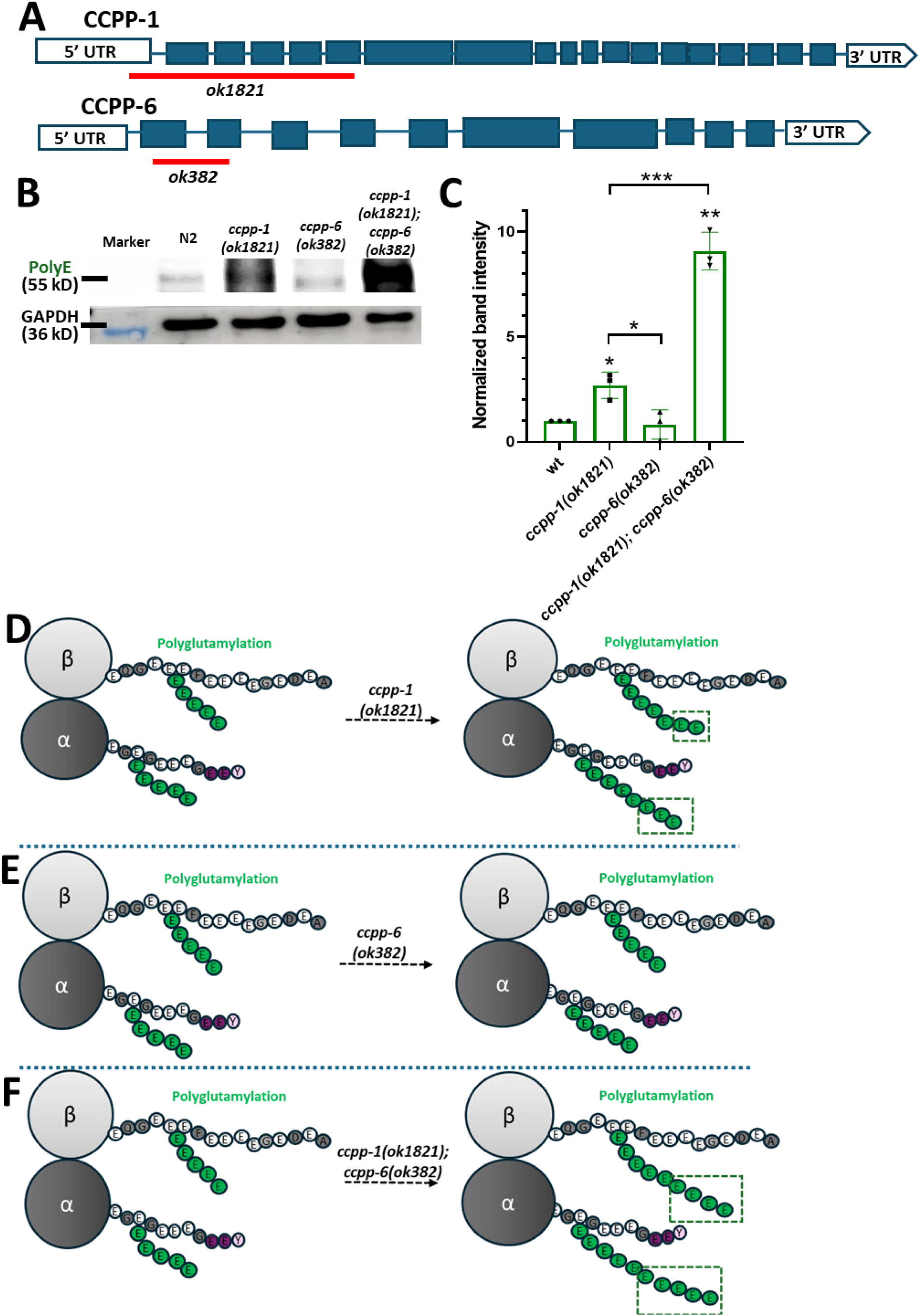
Effect of *ccpp-1* and *ccpp-9* on tubulin polyglutamylation in *C. elegans*. (A) Wild type gene structures of *ccpp-1* and *ccpp-6* as well as available alleles. Red lines indicate deletions in the respective alleles. (B) Western blot employing an antibody detecting polyglutamylated tubulin (PolyE) in N2 wild type, *ccpp-1* and *ccpp-6* single mutants as well as in *ccpp-1;ccpp-6* double mutants. (C) Quantification of data shown in (B). (D-F) Schematic representations of tubulin polyglutamylation effects revealed in (D) *ccpp-1*, (E) *ccpp-6*, and (F) *ccpp-1;ccpp-6* mutants. Dashed green boxes highlight changes in polyglu-tamylation chains. For panel C, one-way ANOVA with Dunnett’s test was used for multiple comparisons with *p<0.05 and **p<0.005. For two-group comparisons, an unpaired, two-tailed Student’s t-test was employed with *p<0.05, **p<0.005 and ***p< 0.001. Box and whisker plots represent the maximum value, upper quartile, median, lower quartile, and minimum value. Experiments carried out in triplicates.

### TTLL-5 but not TTLL-9 acts as a polyglutamylation initiator in *C. elegans*

TTLL enzymes are classified as initiators or elongators based on their enzymatic activity during the polyglutamylation process. In mammals, TTLL4, TTLL5 and TTLL7 initiate polyglutamylation by adding the first glutamate residue to the γ-carboxyl group of a glutamate at the tubulin C-terminal tail, establishing the branching point of the side chain. In contrast, TTLL1, TTLL6, and TTLL11 all function as elongators in mammals, extending that side chain by adding additional glutamate residues (van Dijk et al., 2007). Note that a recent study revealed TTLL11 as an initiator in *Drosophila* (Kravec et al., 2024). To identify the polyglutamylation initiator among the TTLL enzymes in worms, we utilized two available mutant strains: OC343 *ttll-5*(*tm3360*) and OC419 *ttll-9*(*tm3889*). The *ttll-5*(*tm3360*) allele encompasses a 833 bp deletion and a 10 bp insertion, while the *ttll-9*(*tm3889*) allele carries a 509 bp deletion along with a 43 bp insertion. Both mutations introduce frameshifts predicted to produce non-functional proteins due to premature stop codons (Fig. 2A). Western blot analyses revealed that in *ttll-5* mutants, the levels of polyglutamylated tubulin were significantly reduced compared to wild type control, as expected (Fig. 2B-D). In contrast, in *ttll-9* mutants, polyglutamylation levels did not significantly change (Fig. 2B+C, E). Generation of *ttll-5*/*ttll-9* double mutants was not feasible because both alleles are located on the same chromosome V, significantly complicating double mutant isolation due to linkage. Also, multiple attempts to generate the double mutant using CRISPR/Cas9-mediated genome editing were unsuccessful. Nevertheless, these results suggest that in *C. elegans*, TTLL-9 is not essential for initiating polyglutamate chains, while TTLL-5 plays a major role in initiating polyglutamylation.

**Figure 2:**
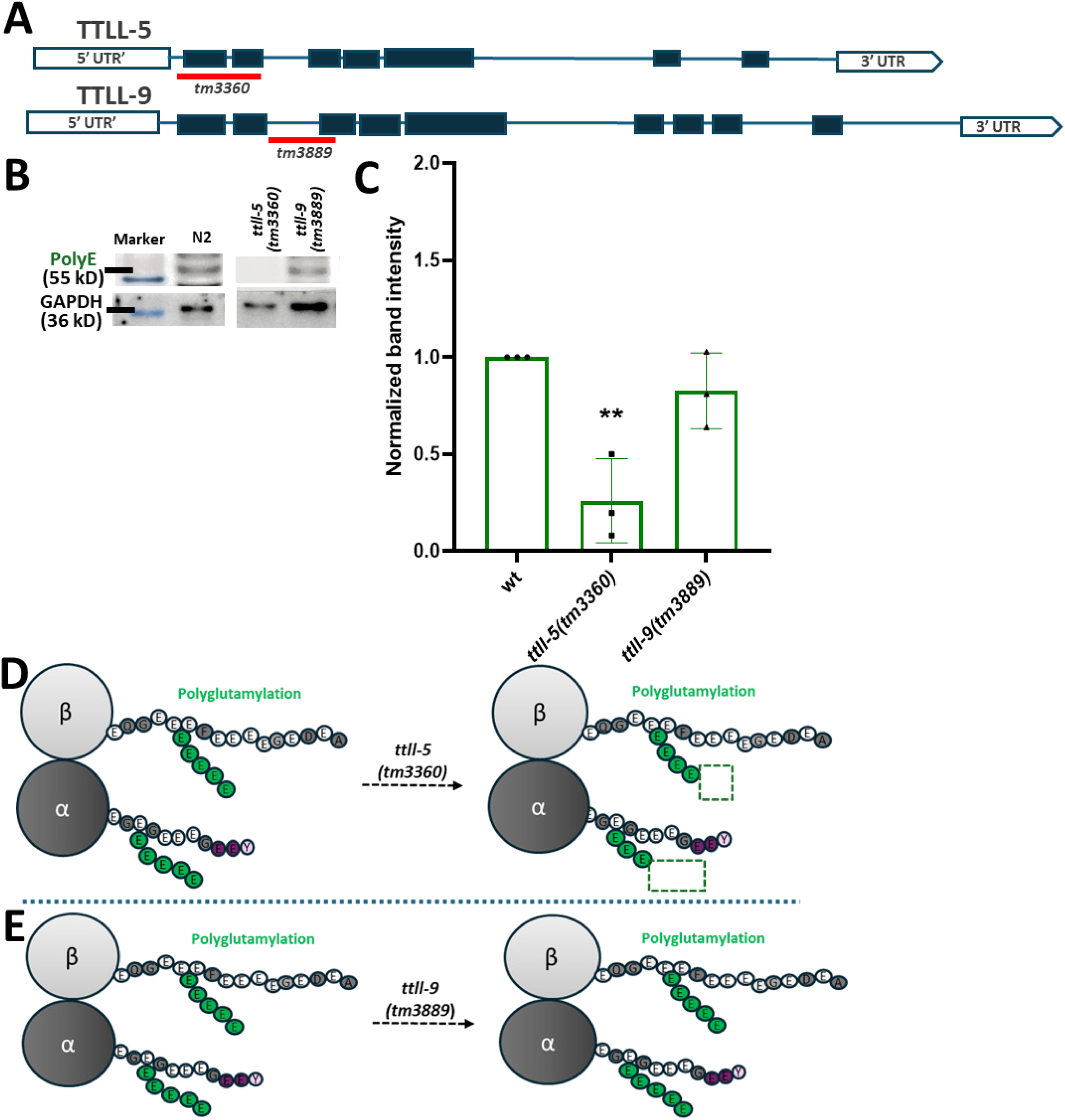
Effect of *ttll-5* and *ttll-9* on tubulin deglutamylation in *C. elegans*. (A) Gene structures of *ttll-5* and *ttll-9* with deletions in mutant alleles highlighted in red. (B) Western blots revealing levels of polyglutamylated tubulin in N2 wild type, *ttll-5* and *ttll-9* mutants. (C) Quantification of Western blot data shown in (B). (D+E) Schematic representations of revealed results for polyglutamylated tubulin in (D) *ttll-5* (D) and (E) *ttll-9*. Dashed green boxes indicate polyglutamylated tubulin modifications. For panel C, oneway ANOVA with Dunnett’s test was used for multiple comparisons with **p<0.005. Box and whisker plots represent the maximum value, upper quartile, median, lower quartile, and minimum value. Experiments replicated in triplicates.

### Crosstalk between tubulin glutamylation and tyrosination

Microtubules are subjected to a variety of PTMs regulating their stability and interactions with MAPs and motor proteins (Janke and Magiera, 2020). Emerging evidence suggests complex crosstalk between polyglutamylation, detyrosination, and acetylation, indicating that these modifications do not work independently but rather influence each other’s dynamics and cellular outcome (Ebberink et al., 2023; Genova et al., 2023; Mahalingan et al., 2020). To explore the interrelationship between these PTMs, we examined the extent of tyrosinated, detyrosinated, and acetylated tubulin in both deglutamylase (*ccpp-1* and *ccpp-6*) and polyglutamylase (*ttll-5* and *ttll-9*) mutants. Western blot analysis revealed that only in *ccpp-1* mutants tyrosinated tubulin levels were significantly elevated; however, remained unchanged in both *ccpp-6* (single) or in *ccpp-1*;*ccpp-6* (double) mutants (Fig. 3A+B, E-G). These data suggest that CCPP-6 acts antagonistically or in a compensatory pathway that suppresses the *ccpp-1* mutant phenotype. In contrast, detyrosinated tubulin levels remain unchanged across single and double mutants (Fig. 3C-G), suggesting a selective effect of deglutamylation on tyrosination but not detyrosination. Intriguingly, tyrosinated tubulin levels were significantly decreased in both *ttll-5*(*tm3360*) and *ttll-9*(*tm3889*) mutants (Fig. 4A+B, E+F), whereas detyrosinated tubulin levels remain unchanged in these strains (Fig. 4C-F). These data reveal a regulatory link in which induced polyglutamylation (*ccpp-1* mutants) promotes tyrosination of tubulin, whereas induced deglutamylation (*ttll-5* mutants) leads to tyrosination reduction. These results can be interpreted by either the existence of not yet characterized tyrosination enzymes in nematodes, or that the glutamylation status of tubulin directly modulates epitopes recognized by the tyrosination antibodies.

**Figure 3:**
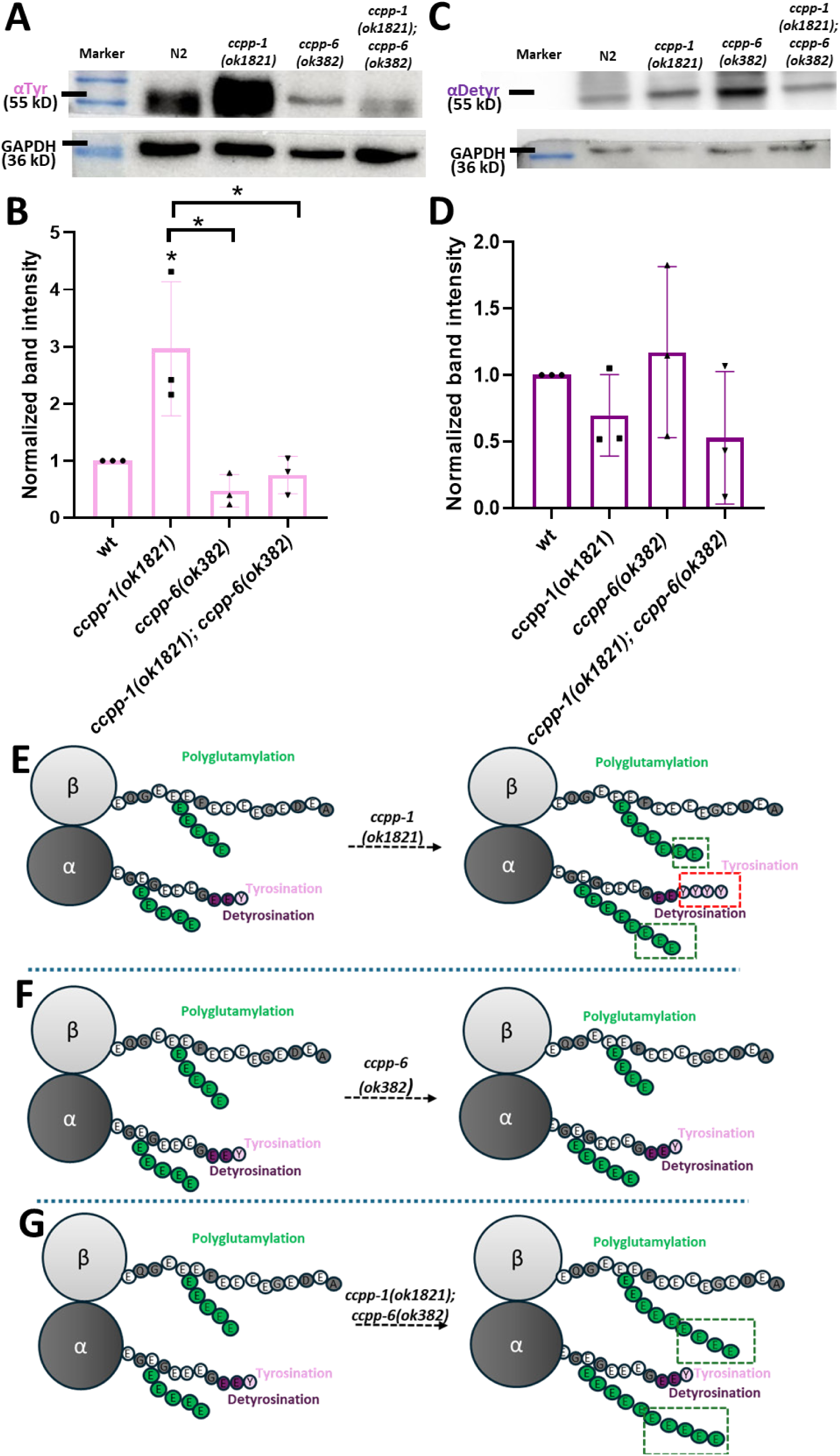
Changes in tubulin tyro– and detyrosination in *ccpp* single and double mutants: (A+C) Western blots displaying signals of (A) anti-tyrosinated tubulin and (C) anti-detyrosinated tubulin in deglutamylase single and double mutants. (B+D) Quantification of data from (A) and (C). (E-G) Schematic representations of changes in tyro– and detyrosinated tubulin in *ccpp* mutants. Dashed green boxes highlight changes in tubulin polyglutamylation and the dashed red box highlights changes in tubulin tyrosination. For panels B+D, one-way ANOVA with Dunnett’s test was used for multiple comparisons (*p<0.05). For two-group comparisons, an unpaired, two-tailed Student’s t-test was employed (*p<0.05). Box and whisker plots represent the maximum value, upper quartile, median, lower quartile, and minimum value. Each experiment was performed in triplicates.

**Figure 4:**
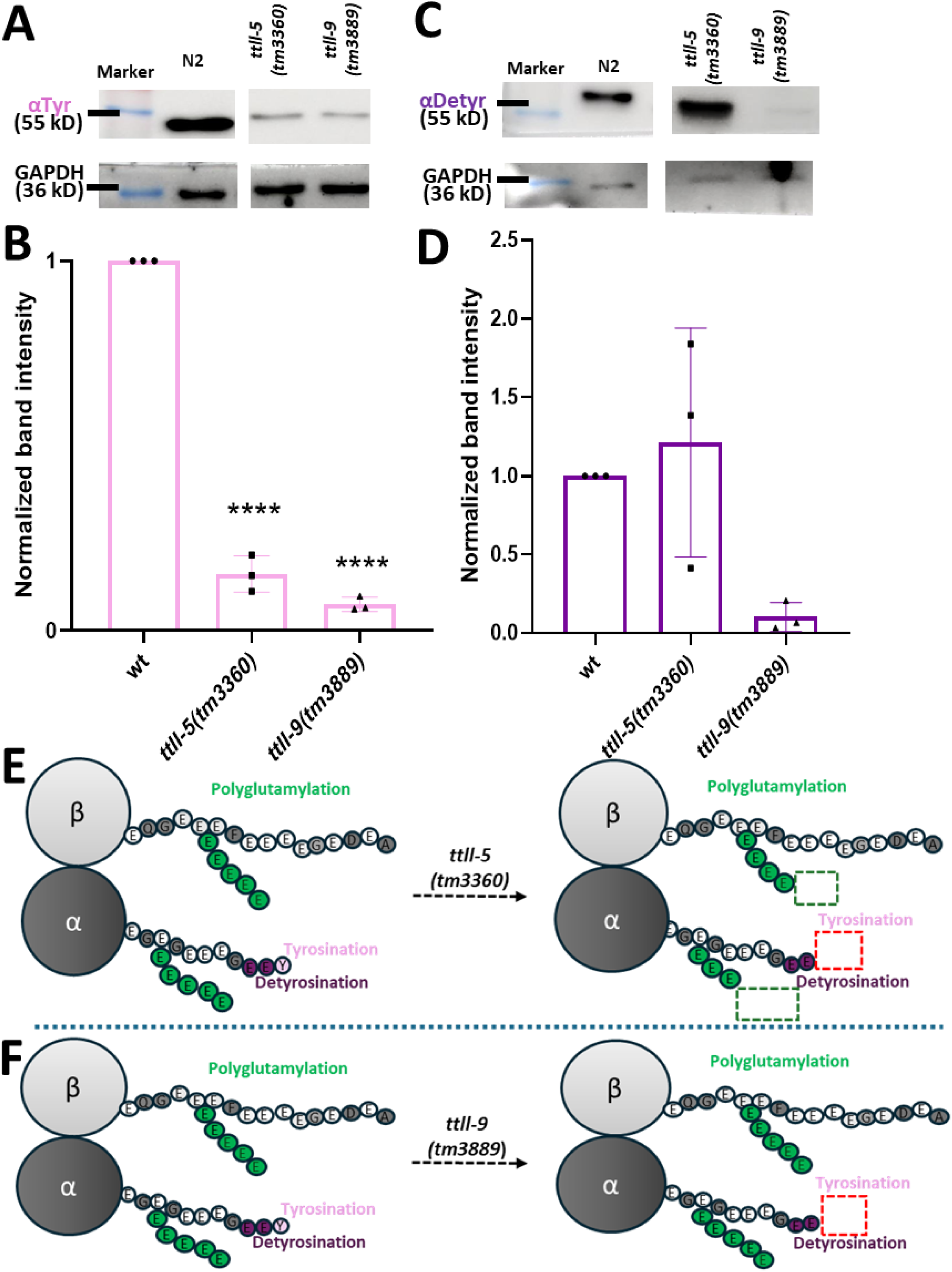
Changes of tubulin tyro– and detyrosination in *ttll-5* and *ttll-9* mutants: (A+C) Western blots showing signals of (A) anti-tyrosinated tubulin and (C) anti-detyrosinated tubulin in *ttll-5(tm3360)* and *ttll-9(tm3889)* mutants. (B+D) Quantification of data from (A) and (C). (E+F) Schematic representations of changes in tyro– and detyrosinated tubulin in *ttll-5* and *ttll-9* mutants, respectively. Dashed green boxes highlight changes tubulin polyglutamylation and the dashed red box highlights changes in tubulin tyrosination. For panels B+D, one-way ANOVA with Dunnett’s test was used for multiple comparisons with ****p<0.0001. Box and whisker plots represent the maximum value, upper quartile, median, lower quartile, and minimum value. Each experiment was performed in triplicate.

To further explore the relationship between polyglutamylation and other PTMs, we assessed levels of acetylated tubulin. However, no significant changes were observed in either deglutamylase or polyglutamylase (*ttll-5*) mutants (Fig. 5), suggesting no functional crosstalk between glutamylation and acetylation. Together, these findings indicate that glutamylation selectively regulates the tyrosination state of tubulin, but not its acetylation or detyrosination. This highlights an important functional tubulin PTM crosstalk that fine-tunes microtubule dynamics and their interactions with MAPs and motor proteins.

**Figure 5:**
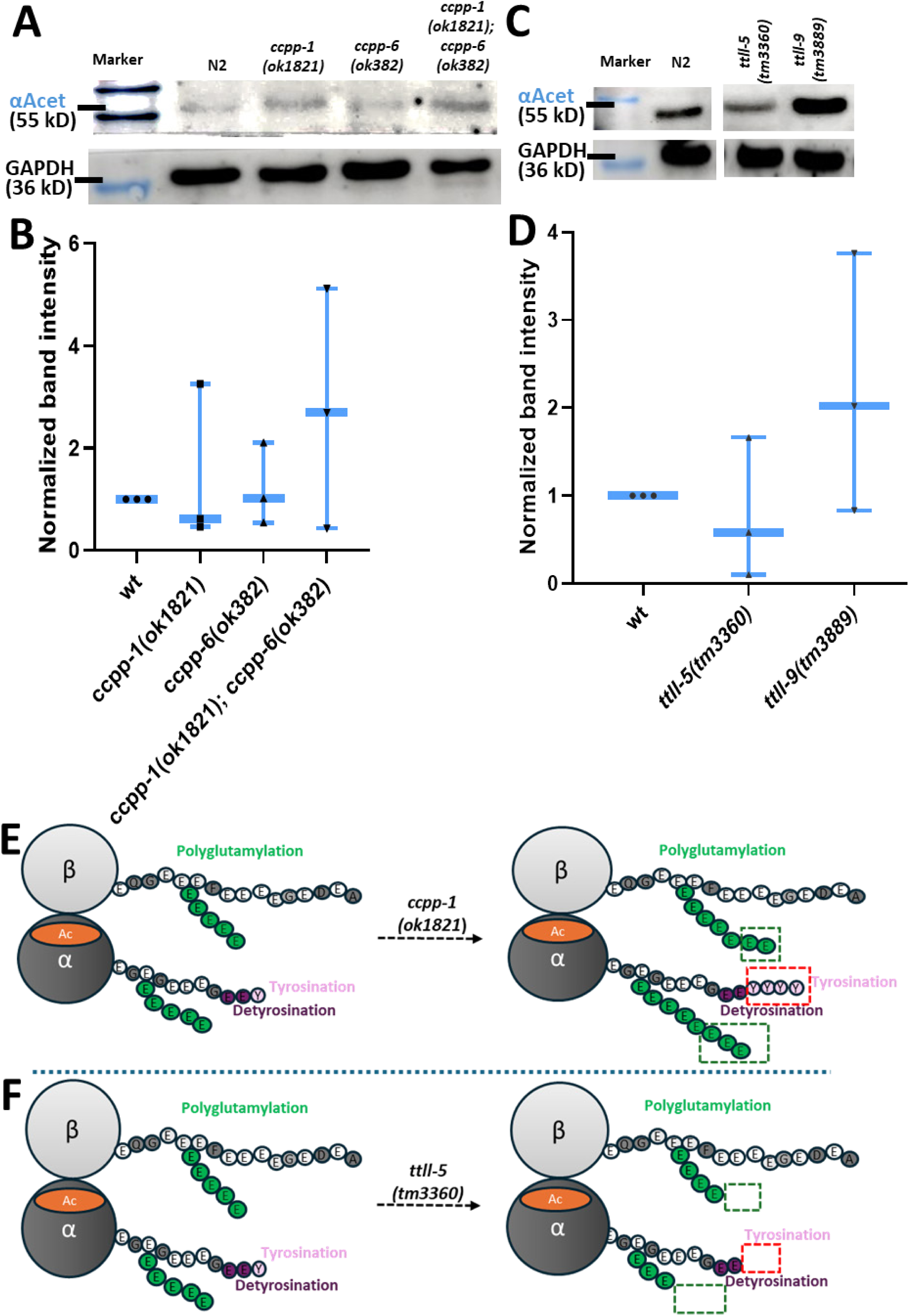
Effects of tubulin acetylation in deglutamylase and polyglutamylase mutants: (A+C) Western blots displaying signals of α-acetylated tubulin levels in (A) deglutamyalse mutants and (C) polyglutamylase mutants. (B+D) Quantification of data from (A) and (C). (E+F) Schematic representations of changes in acetylated tubulin in *ccpp-1(ok1821)* and *ttll-5(tm3360)* mutants. Dashed green boxes highlight changes in tubulin polyglutamylation and the dashed red box highlights changes in tubulin tyrosination. Orange circles depict acetylated tubulin. For panels B+D, one-way ANOVA with Dunnett’s test was used for multiple comparisons. Box and whisker plots represent the maximum value, upper quartile, median, lower quartile, and minimum value. Each experiment was performed in triplicate.

### Hyperglutamylated tubulin negatively affects kinesin-3 transport

To determine the functional consequences of this PTM crosstalk, we assessed the motility of the most important axonal motor protein UNC-104, tagged with monomeric red fluorescent protein (mRFP). This fluorescent tag was selected for its low self-aggregation properties, allowing visualization of motor movement in live animals without artificial fluorophore aggregation (Eves-van den Akker et al., 2014; Wang et al., 2019). We specifically employ the ALM (*a*nterior *l*ateral *m*icrotubule cell) neuron, spreading nearly 50% of the worm’s body, allowing for long-range observations of motility (Suppl. Fig. S1A). Figure 6A exhibits that anterograde velocities of UNC-104 are significantly reduced in *ccpp-1* and *ccpp-6* mutants and that this effect is further enhanced in double mutants (pointing to parallel – and partially redundant – pathways). While anterograde run lengths of UNC-104 remain unchanged in single mutants, significant changes occur in double mutants, consistent with a synthetic phenotype (Fig. 6B). Though UNC-104 is an anterograde motor and does not inherit the ability to move retrogradely, it is interesting to further analyze its retrograde features (likely based on direct interactions with the retrograde motor dynein). Indeed, direct interactions between UNC-104 and dynein have been reported (Chen et al., 2019), often leading to tug-of-war events and eventually passive retrograde displacements of UNC-104 (Tien et al., 2011; Wagner et al., 2009). Interestingly, retrograde displacements of UNC-104 follow a similar pattern observed for anterograde movements, e.g., if anterograde velocities or run lengths are diminished in mutants, they are also diminished in retrograde directions (Fig. 6A-D). The “paradox of co-dependence model” may explain this finding: motors are mutually activated by mechanically pulling on each other in opposing directions. If this mechanical activation is lost or reduced, motor activation is diminished (Hancock, 2014). Interestingly, for these retrograde features of UNC-104, the double mutant mostly phenocopies *ccpp-6* (Fig. 6C+D), leading to the assumption that linear epistasis exists with CCPP-6 being downstream of CCPP-1. We then determined the number of directional changes of motors (as a measure of tug-of-war intensity). A significant increase in these motor reversals was quantified in *ccpp-6* and *ccpp-1/ccpp-6* double mutants (Fig. 6E). A significant increase in motor pausing was observed in double mutants as well (Fig. 6F), consistent with the strong reduction in velocities in these mutants (Fig. 6A); However, moving persistence at uniform velocities (PMUV) did not significantly change across the mutants (Fig. 6G).

**Figure 6:**
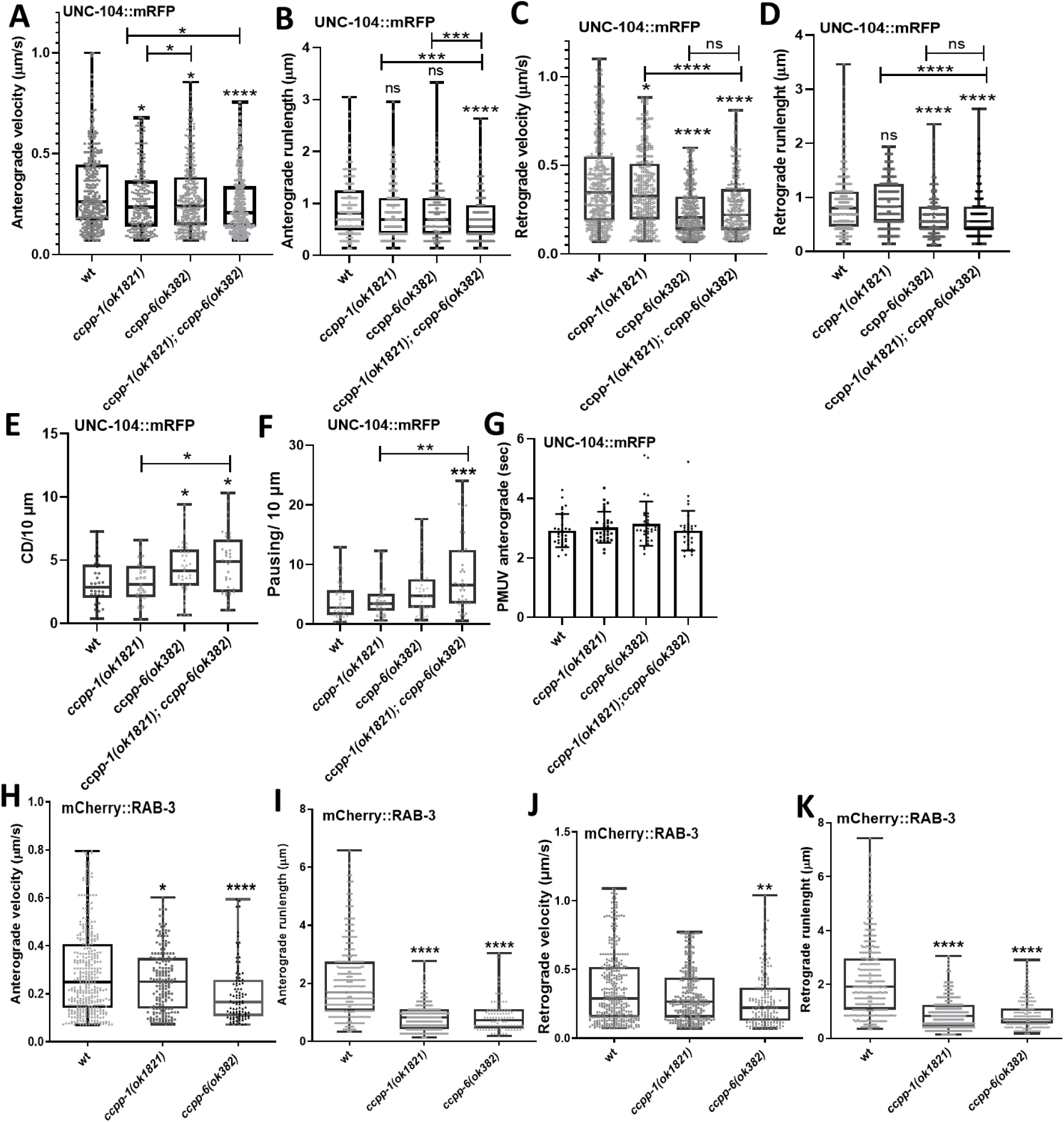
Hyperglutamylated tubulin negatively affects kinesin-3-based axonal transport. (A) Anterograde velocity of UNC-104::mRFP in wt, *ccpp-1, ccpp-6* as well as in *ccpp-1;ccpp-6* double mutants. (B) UNC-104 anterograde event run lengths. (C) UNC-104 retrograde velocity and (D) retrograde event run lengths. (E) Changes of directions of UNC-104 within 10 µm. (F) Number of UNC-104 pauses per 10 µm range. (G) Persistent motility at uniform velocities in anterograde direction. Note that “wild type” refers to a strain that has the uncoordinated phenotype – caused by a point mutation in the unc-104 gene (*e1265* allele) – rescued by pUNC-104::UNC-104::mRFP expression. (H) mCherry::RAB-3 anterograde velocity. (I) RAB-3 anterograde run length. (J) RAB-3 retrograde velocity. (K) RAB-3 retrograde run length. Analyzed total events for A-G: wt = 1009, *ccpp-1*(*ok1821*) = 781, *ccpp-6*(*ok382*) = 1108, *ccpp-1*(*ok1821*)*; ccpp-6*(*ok382*) = 1028. Analyzed total events for H-K: wt = 798, *ccpp-1(ok1821*) = 652 and *ccpp-6(ok382)* = 412. One-way ANOVA with Dunnett’s test with *p<0.05, **p<0.005 and **** p<0.0001 in (A+C+H+J). Non-parametric Kruskal-Wallis one-way ANOVA with Dunn’s test for multiple comparisons with *p<0.05, **p<0.005, ***p< 0.001, and ****p<0.0001 in (B, D-G, I+K). For two group comparisons in (A) unpaired Student’s t-test with *p<0.05 was used. For (B, D-F) we use unpaired, non-parametric Mann-Whitney test with *p<0.05, **p<0.01, ***p<0.001. Box and whisker graphs represent maximum value, upper quartile, median, lower quartile and minimum value.

We then examined the motility of UNC-104’s major cargo RAB-3 tagged with mCherry (another monomeric fluorophore designed to exhibit reduced aggregation propensities). It is expected that the motility of the major cargo alone should be comparable to the motility of its major transporter (UNC-104) alone. Indeed, loss of CCPP-1 and CCPP-6 significantly reduces both anterograde velocities as well as run lengths of RAB-3 (Fig. 6H+I). Similarly, retrograde velocity (except for *ccpp-1*) and run lengths of RAB-3 were significantly decreased (Fig. 6J+K). These results suggest that hyperglutamylation (and tyrosination increase via crosstalk) perturbs the entire motor/cargo transport complex.

Along with decreased moving qualities of UNC-104 upon hyperglutamylation (Fig. 6), motors are visibly retained in cell bodies of *ccpp-1* and *ccpp-1/ccpp-6* double mutants (Fig. 7A+B, cell body) with an overall reduced presence in axons (Fig. 7A+B, initial, middle and distal region). Additionally, travel distances (from axon hillocks to distal regions) are reduced as well (Fig. 7C, Suppl. Fig. S1D). We then wondered if correlations exist between motor speeds (Fig. 6A) and travel distances (Fig. 7C). Indeed, strong positive correlations are determined between motor speeds and travel distances across all single and double deglutamylase mutants (Fig. 7D-G).

**Figure 7:**
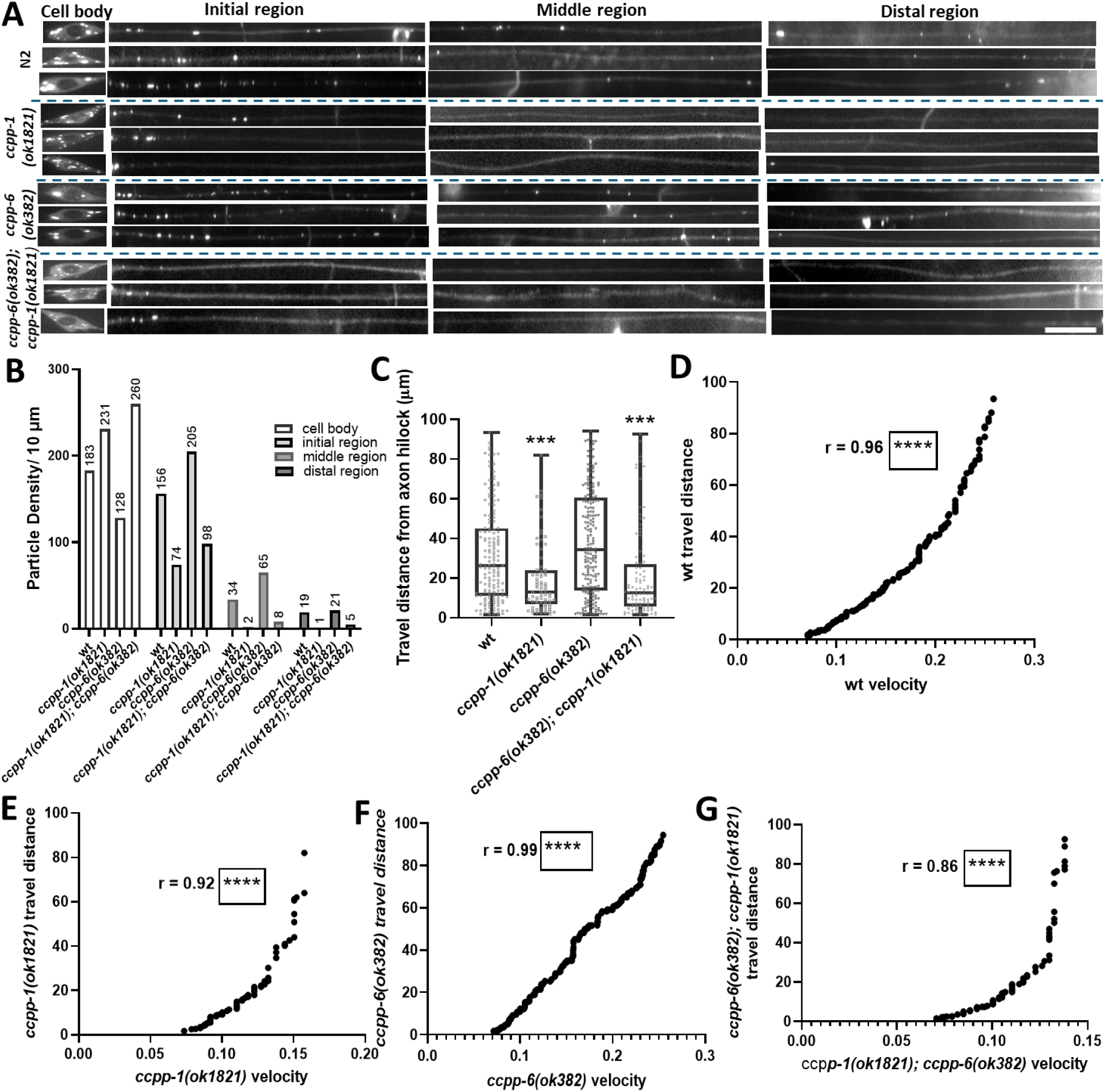
Effect of deglutamylase enzymes on UNC-104 axonal cluster patterns. (A) Stacks of ALM neurons (after digital straightening) to examine UNC-104 cluster patterns in single and double deglutamylase mutants. (B) Density of UNC-104 particles in cell body, initial, middle and distal regions. (C) Travel distances of UNC-104 (data taken from Fig. 7A). (D-G) Correlation between UNC-104 motility (taken from Fig. 6A) and UNC-104 particle travel distance in ALM neurons (taken from Fig. 7C) in (D) wild type, (E) *ccpp-1*, (F) *ccpp-6* and (G) *ccpp-1*(*ok1821*)*;ccpp-6*(*ok382*) double mutants. “Wild type” refers to the pUNC-104::UNC-104::mRFP(*e1265*) rescue strain. N = 20-25 worms per experimental group for cluster analysis in (A-C). Non-parametric Kruskal-Wallis one-way ANOVA with Dunn’s test for multiple comparisons with ***p< 0.001 in (C). R values based on Pearson’s correlation coefficient and p values determined using Student’s t-test for two group comparisons with ****p<0.0001. Box and whisker graphs represent maximum value, upper quartile, median, lower quartile and minimum value. Scale bar: 10 µm.

### Hypoglutamylated tubulin positively affects UNC-104 transport

Since hyperglutamylated tubulin negatively affects the motility of UNC-104, we then investigated the effect of hypoglutamylation on axonal transport. In *ttll-5(tm3360)* as well as *ttll-9(tm3889)* mutants, both anterograde velocities and run lengths of UNC-104 were significantly increased (Fig. 8A+B). In addition, moving persistence increased as well in both polyglutamylase mutants (Fig. 8C). These results reveal an enhancement of travel qualities of UNC-104 upon reduced tubulin glutamylation (*ttll-5*, Fig. 2) and reduced tyrosination (*ttll-5* and *ttll-9*, Fig. 4). These improved motor movements are directly reflected in a reduction of motors in neuronal cell bodies (Fig. 8D+E, cell body) and a higher presence in axons (Fig. 8D+E, initial, middle and distal region). Also travel distances are significantly increased in *ttll-9* mutants (though for *ttll-5* this parameter remains unchanged) (Fig. 8F). Because *ttll-9* reduces tyrosination (Fig. 4B) and not polyglutamylation (Fig. 2C), we assume that positively affected travel distances are a result of more stable microtubules. From these results, we conclude that a reduction in polyglutamylation (as well as tyrosination reduction via cross-talk) enhances UNC-104 motility.

**Figure 8:**
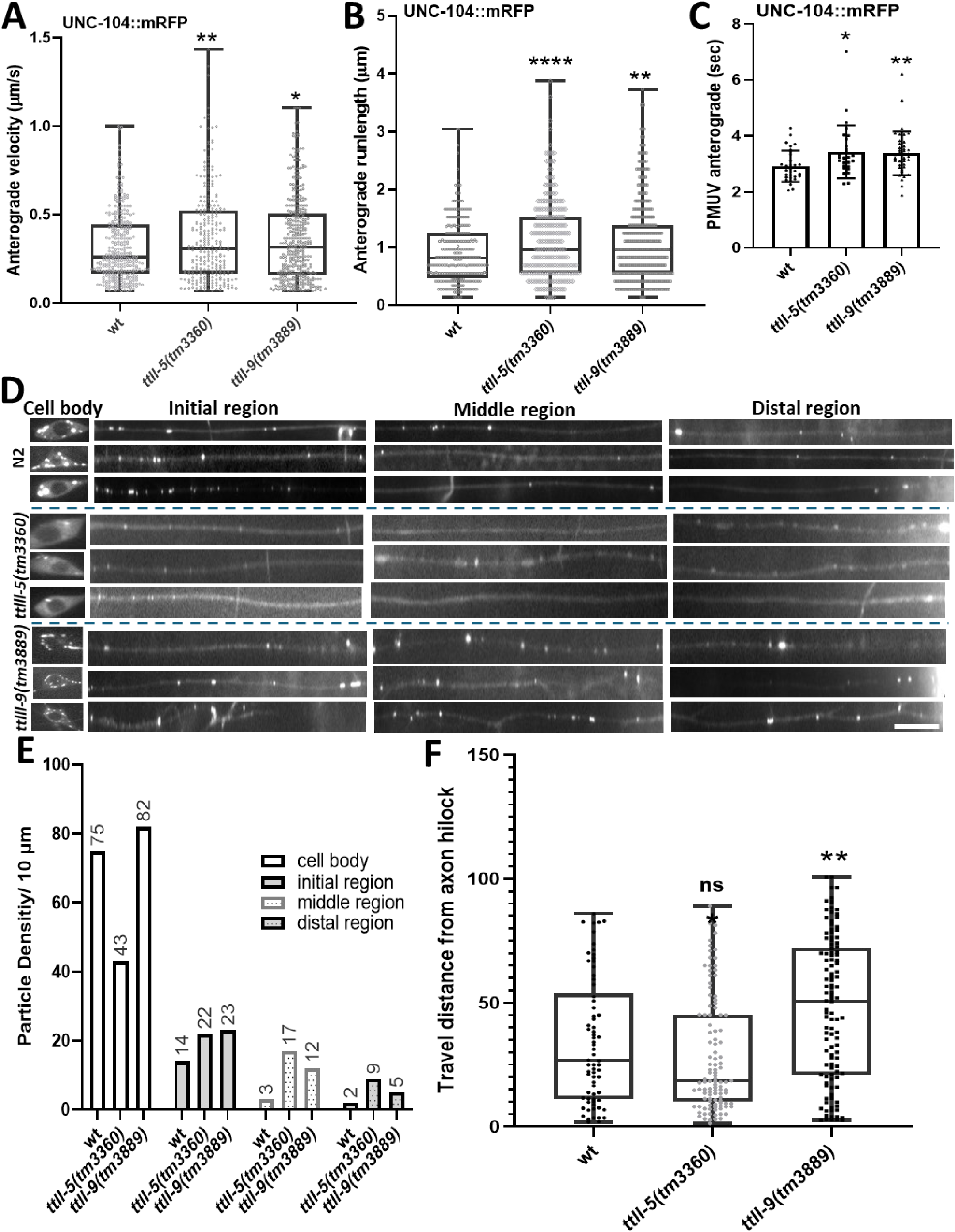
Effect of hypoglutamylated tubulin on UNC-104 motility. (A) Anterograde velocity of UNC-104::mRFP in wt, *ttll-5* and *ttll-9* mutants. (B) UNC-104 anterograde event run lengths. (C) Persistent motility at uniform velocities in anterograde directions. (D) Stacks of ALM neurons in wild type, *ttll-5* and *ttll-9* mutants. (E) Density of UNC-104 particles per 10 µm in cell body, initial, middle and distal regions. (F) Travel distances of UNC-104 (data taken from Fig. 8D). Analyzed total events: wt = 1009, *ttll-5*(*tm3360*) = 710 and *ttll-9*(*tm3889*) = 1143. One-way ANOVA with Dunnett’s test with *p<0.05 and **p<0.005 in (A). n = 11 worms for wt, *ttll-5(tm3360)* and *ttll-9(tm3889)* mutants in (D). Note that for this Figure “wild type” refers to the pUNC-104::UNC-104::mRFP(*e1265*) rescue strain. Non-parametric Kruskal-Wallis one-way ANOVA with Dunn’s test for multiple comparisons with *p<0.05, **p<0.005 and ****p<0.0001 in (B+C, F). Box and whisker graphs represent maximum value, upper quartile, median, lower quartile and minimum value. Scale bar: 10 µm.

### Colocalization between glutamylase enzymes and UNC-104

Above, we validated the functional effects of CCPP and TTLL enzymes on the quality of axonal transport in *C. elegans*. To understand whether these enzymes indeed co-express with the molecular motor UNC-104 in neurons, we generated transcriptional reporter fusions of ccpp-1, ccpp-6 and ttll-5 and expressed them along with pUNC-104::UNC-104::mRFP in worms. From Figure 9A, it is evident that pCCPP-1::GFP is clearly expressed in various nerve ring neurons, ALM, tail neurons as well as in male-specific sensory neurons. UNC-104 significantly colocalizes with pCCPP-1::GFP as quantified by Mander’s and Pearson’s coefficients as well as PDM images (Fig. 9A-D). Though pCCPP-6::GFP was overall expressed more weakly, we additionally identified expression in sublaterals as well as in dorsal axons (Fig. 9B). Colocalization with UNC-104 was present though weaker as well (likely as a result of the weaker CCPP-6 expression) (Fig. 9B, E+F). We also examined the expression pattern of a transcriptional TTLL-5 fusion (Fig. 9G) and identified expression in head sensory neurons, dorsal and ventral nerve cords, as well as in the epidermis. Also for this enzyme, colocalization with UNC-104 is present, however, rather weak (Fig. 9G, H+I). We then conducted whole-mount immunostaining employing pUNC-104::UNC-104::GFP(*e1265*) rescue worms and staining them with anti-polyglutamylated tubulin as well as anti-tyrosinated tubulin antibodies. Though we found whole-mount immunostainings of worms technically challenging, we were able to receive antibody signals from polyglutamylated tubulin as well as tyrosinated tubulin colocalized with expressed UNC-104::mRFP (Suppl. Fig. S2C. Specifically, tyrosinated tubulin revealed distinct colocalization patterns (Suppl. Fig. S2C).

**Figure 9:**
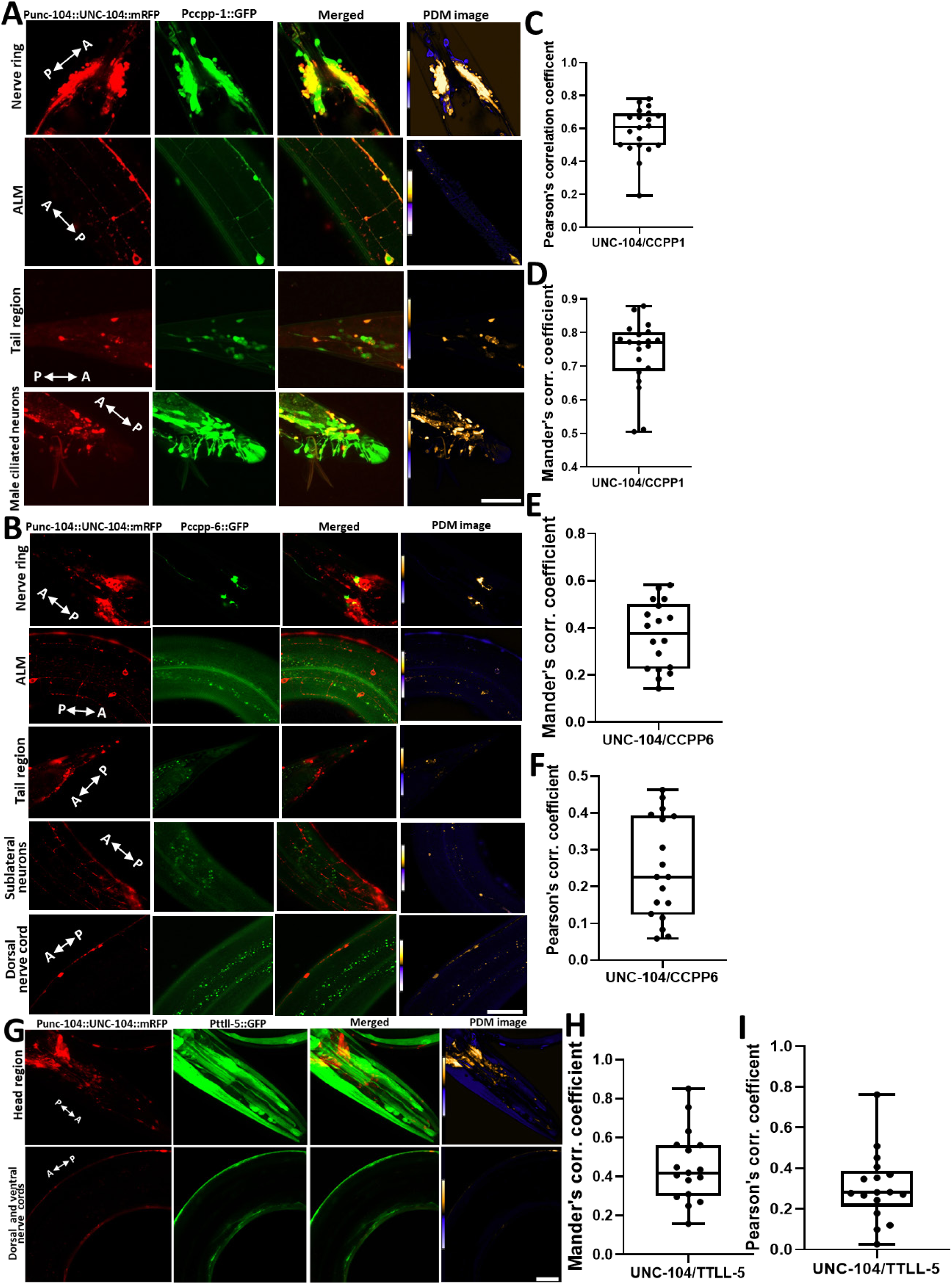
Colocalization between UNC-104 and transcriptional ccpp-1, ccpp-6 and ttll-5 fusions. (A) Colocalization between UNC-104 and ccpp-1 promoter fusions in *C. elegans* neurons. (C+D) Mander’s and Pearson’s coefficients from data shown in (A). (B) Colocalization between UNC-104 and ccpp-6 promoter fusions in *C. elegans* neurons. (E+F) Mander’s and Pearson’s coefficients from data shown in (B). (G) Colocalization between UNC-104 and ttll-5 promoter fusion in *C. elegans* neurons. (H+I) Mander’s and Pearson’s coefficients from data represented in (G). Box and whisker graphs represent maximum value, upper quartile, median, lower quartile and minimum value. N = 18-20 worms. A = anterior, P = posterior directions. Scale bar: 10 µm.

### Polyglutamylated tubulin associates with kinesin-3

We then challenged the possibility of physical associations between polyglutamylated tubulin and UNC-104 and whether such interactions would change in *ccpp-1* and *ttll-5* mutants. Here, we performed co-immunoprecipitation (Co-IP) assays to first precipitate UNC-104 (using a commissioned anti-UNC-104 antibody) and then to detect PolyE tubulin employing lysates of worms carrying *ttll-5* and *ccpp-1* alleles. As a result, in *ttll-5(tm3360)* mutants (with significantly reduced polyglutamylated tubulin levels, Fig. 2B+C), we observed reduced binding of PolyE tubulin to UNC-104 (Fig. 10A-D). Conversely, in *ccpp-1(ok1821)* mutants (with significantly increased polyglutamylated tubulin levels, Fig. 1B+C), polyglutamylated tubulin binding to UNC-104 significantly increased (Fig. 10E–G). These findings demonstrate that polyglutamylated tubulin physically associates with UNC-104 and that the presence or absence of glutamylation enzymes significantly regulates this interaction.

**Figure 10:**
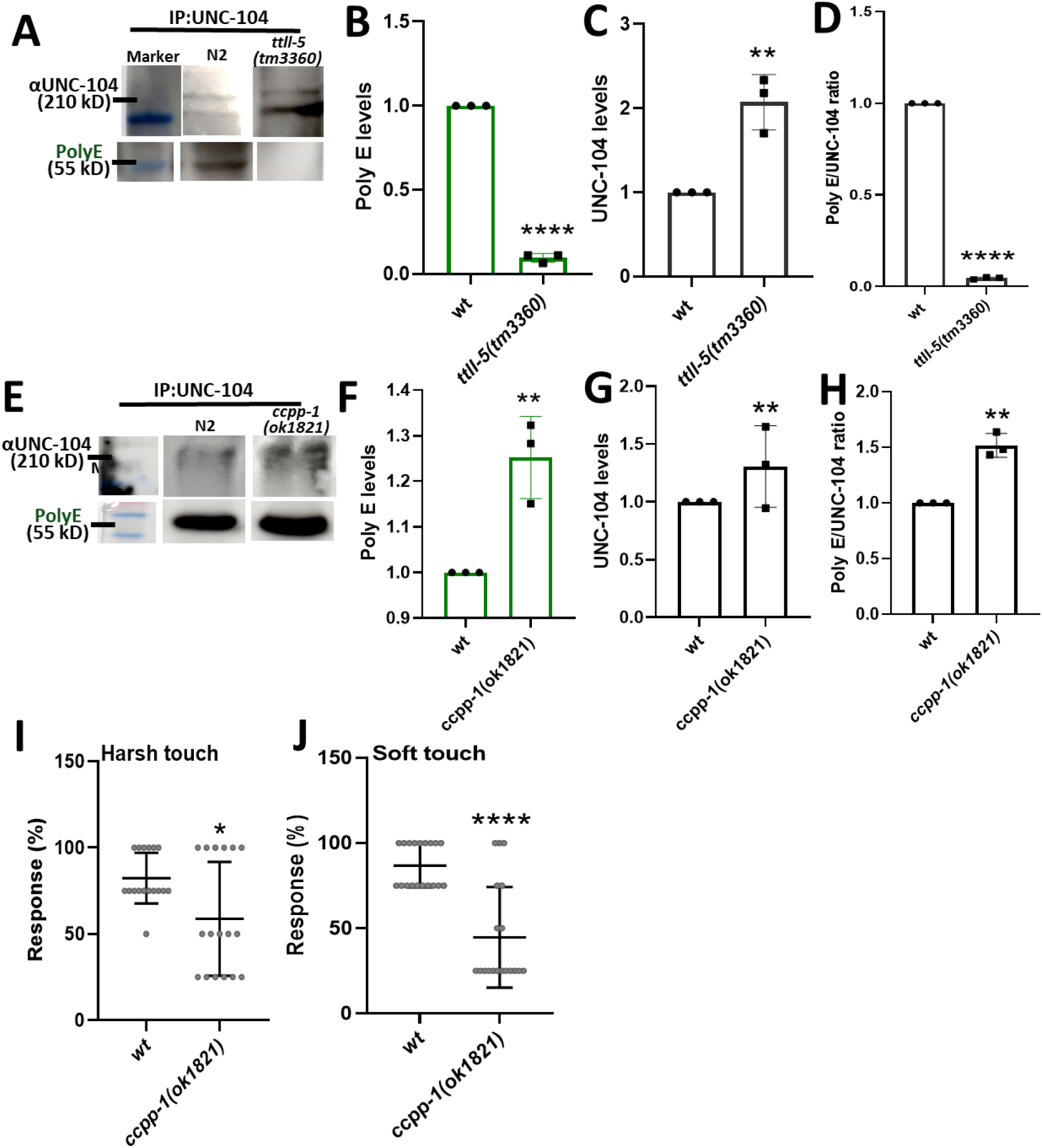
Polyglutamylated tubulin physically associates with UNC-104. (A+E) Co-immunoprecipitation assays showing the interaction between UNC-104 and polyglutamylated tubulin using lysates from whole worms expressing UNC-104::mRFP in (A) wt and polyglutamylase *ttll-5*(*tm3360*) mutants and (E) wt and deglutamylase *ccpp-1*(*ok1821*) mutants. (B-D) Co-IP quantification of (B) polyglutamylated tubulin, (C) UNC-104 protein and (D) PolyE/UNC-104 ratio in wt and *ttll-5*(*tm3360*) mutants. (F-H) Co-IP quantification of (F) polyglutamylated tubulin, (G) UNC-104 protein and (H) PolyE/UNC-104 ratio in wt and *ccpp-1*(*ok1821*) mutants. Unpaired, two-tailed Student’s t-test with **p<0.005. Number of trials for each experimental group: 3. (I) Harsh– and (J) soft touch assay in *ccpp-1(ok1821)* mutants. Unpaired, non-parametric Mann-Whitney test with *p<0.05, **p<0.01, ***p<0.001 and ****p<0.0001. n = 13-19 worms.

### Physiological consequences of reduced axonal transport efficiency in *ccpp-1* mutants

To investigate whether reduced transport efficiency in *ccpp-1* mutants would have any physiological consequences for the worms, we performed behavioral assays measuring responses to both gentle (soft) and harsh touch stimuli, which are mediated by mechanosensory neurons such as ALM. Consistent with disrupted neuronal function, *ccpp-1* mutants displayed significantly diminished responses in both touch assays compared to wild-type animals (Fig. 10I+J). These findings indicate that impaired tubulin glutamylation and its effects on axonal transport negatively impact mechanosensory behavior. Therefore, *C. elegans* can serve as a useful model organism for studying the role of tubulin glutamylation and associated neuropathological defects.

### Model summarizing this study

In our model (Fig. 11), we propose that increased polyglutamylation physically “traps” kinesin-3 motors along the microtubule based on charge interactions: polyglutamylated tubulin is strongly negatively charged (Janke et al., 2008) that increasingly interacts with the positively charged K-Loop in the motor domain of UNC-104 (Fig. 11B) (Benoit et al., 2024; Soppina and Verhey, 2014) leading to UNC-104 immobilization. However, for microtubules with hypoglutamylated tubulin (Fig. 11C) this tubulin “stickiness” is reduced, and motors can move more freely. In addition, increased hyperglutamylation is linked to elevated microtubule severing (Lacroix et al., 2010; Valenstein and Roll-Mecak, 2016), and increased tyrosination is linked to elevated microtubule depolymerization (Sanyal et al., 2023) (depicted as shorter MTs in Fig. 11), leading to less efficient axonal transport.

**Figure. 11:**
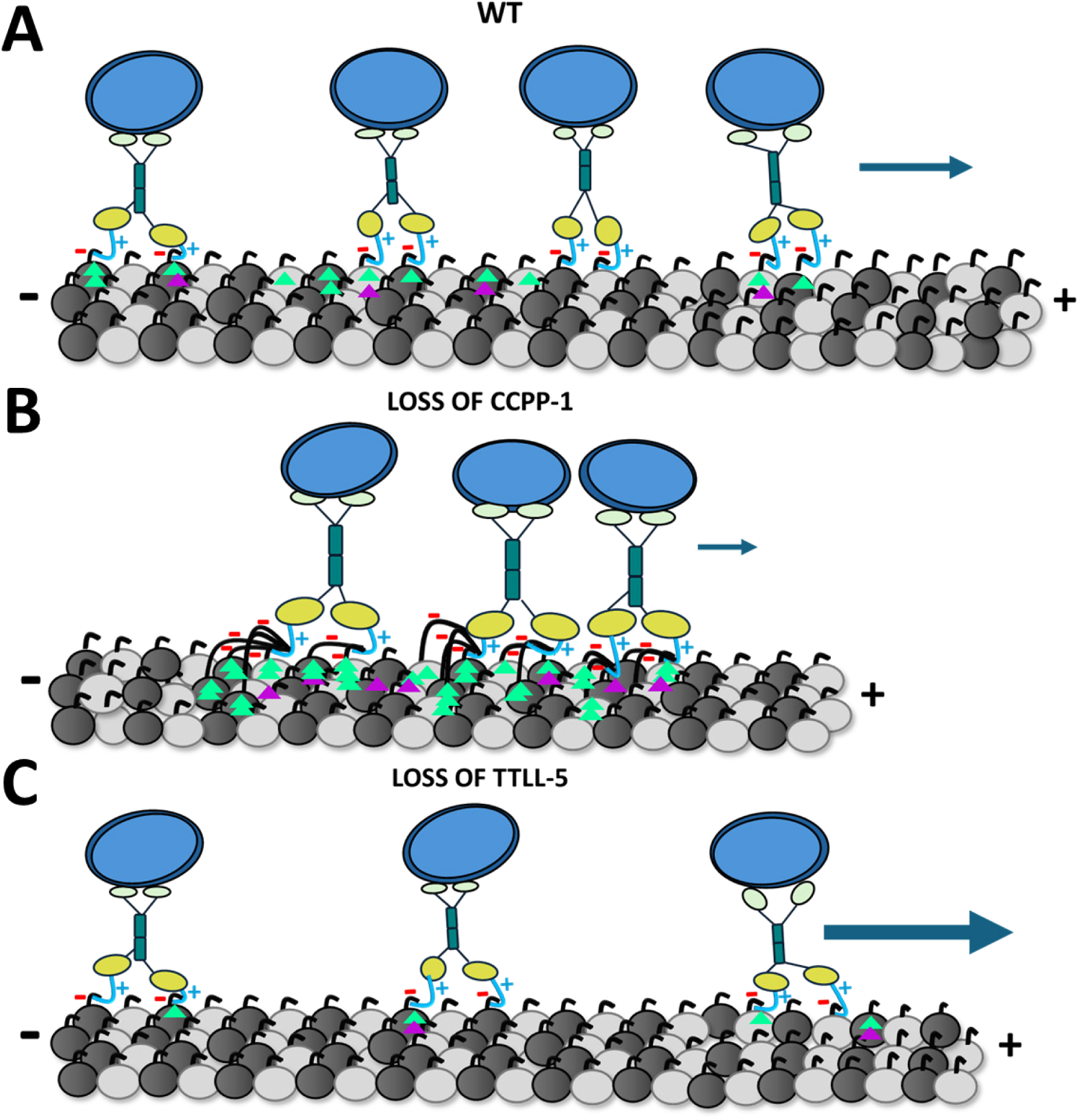
Model summarizing the findings of this study. (A) UNC-104 motors (carrying synaptic vesicles) moving on a microtubule in wild type animals, (B) *ccpp-1* mutants and (C) *ttll-5* mutants. The blue arrow represents relative speeds. Positively charged (blue plus signs) K-loop of UNC-104 (light blue hook) interacts with the negatively charged (red minus signs) C-terminal tail (E-hook, black hook) of tubulin. Green triangles represent polyglutamylation and magenta triangles indicate tyrosination. Black plus– and minus signs represent the polarity of the microtubule. Increased polyglutamylation leads to increased negative surface charges of the microtubule, increasingly trapping and immobilizing UNC-104 in *ccpp-1* mutants. In addition, increased tyrosination and polyglutamylation leads to less stable microtubules (depicted as shorter microtubules) (B). Decreased polyglutamylation leads to motors moving more freely UNC-104 in *ttll-5* mutants (C).

## DISCUSSION

### Genetic pathways of ccpp-1 and ccpp-6 in *C. elegans*

Microtubules are highly dynamic cytoskeletal elements that undergo continuous assembly and disassembly, playing essential roles in intracellular transport, cell polarity and cell shape. These dynamics are influenced by intrinsic factors such as tubulin isoforms, various PTMs, and MAPs, as well as extrinsic factors including microtubule-interfering drugs and temperature (Janke and Magiera, 2020). *C. elegans* possesses only two deglutamylase enzymes: CCPP-1 and CCPP-6. As shown in Fig. 1B+C, the level of polyglutamylated tubulin was elevated in *ccpp-1(ok1821)* mutants and further increased in the *ccpp-1;ccpp-6* double mutants (with no changes in single *ccpp-6(ok382)* backgrounds). An enhanced phenotype of one gene in double mutants (with no phenotype for the other gene) often points to parallel and partially redundant pathways. Consistently, in a study investigating the effects of CCPP-1 and CCPP-6 on *C. elegans* cilia development, authors conclude no redundancy of CCPP-6 (Klimas et al., 2023). In a study using mice, deletion of CCP1 leads to hyperglutamylation in the cerebellum as expected, but led to a *pcd* (Purkinje cell degeneration) mouse phenotype (Fernandez-Gonzalez et al., 2002; Rogowski et al., 2010). On the other hand, loss of CCP6 caused hyperglutamylation only restricted to specific brain regions and did not produce a *pcd* phenotype (Kalinina et al., 2007). Notably, simultaneous deletion of both CCP enzymes induced tissue widespread hyperglutamylation and neuronal degeneration throughout the mouse brain, including in regions that remain unaffected in *pcd* mice, such as the pyramidal neurons of the cerebral cortex (Magiera et al., 2018). This evidence supports that both CCP1 and CCP6 indeed function as deglutamylases, but work in distinct pathways. Redundancy might also explain the differing effects of CCPP-1 and CCPP-6 on UNC-104 cluster density patterns (Fig. 7A+B): motor retention in neuronal cell bodies was more pronounced in *ccpp-1* mutants as compared to *ccpp-6* mutants, and while travel distances were significantly decreased in *ccpp-6* mutants, we could not detect such an effect in *ccpp-6* mutants. In summary, from the literature and our experiments, we conclude partially redundancy for genes ccpp-1 and ccpp-6.

### Hyperglutamylation negatively affects kinesin-3-based axonal transport in nematodes

We demonstrated that in deglutamylase mutants, tubulin is hyperglutamylated as expected (Fig. 1) and that this effect negatively affects UNC-104 motility (Fig. 6A+B) as well as cargo transport efficiencies (Fig. 6H+I). Vice versa, in polyglutamylase mutants with hypoglutamylated tubulin (Fig. 2), motor motility significantly increased (Fig. 8A-C). These findings are consistent with single-molecule *in vitro* assays, indicating that tubulin polyglutamylation decreases KIF1A run length by 34% and landing rates by 94% (Lessard et al., 2019). Likewise, another group found that in *Ccp1*^−/−^*Ccp6*^−/−^ mice (with hyperglutamylated tubulin) kinesin-1-based mitochondria transport is perturbed in neurons leading to neuronal degeneration (Magiera et al., 2018). To explain the impaired UNC-104 motility by hyperglutamylation (Figs. 6, 7), we propose that motors are increasingly immobilized on microtubules due to increased negative surface charges upon tubulin hyperglutamylation. Indeed, it has been well documented that KIF1A binds to microtubules via its positively charged K-loop (in the motor domain) that charge-interacts with the negatively charged C-terminal E-Hooks of β-tubulin (Benoit et al., 2024; Zaniewski and Hancock, 2023). In our model in Figure 11, increased “stickiness” of microtubules due to hyperglutamylation increasingly traps motors on the microtubule surface, thereby slowing transport. It has also been reported that hyperglutamylated tubulin destabilizes microtubules in various ways: promoting binding of microtubule severing enzymes, disrupting MAP binding and increasing microtubule catastrophe (Chen and Roll-Mecak, 2023). In more detail, long polyglutamylated side chains enhance spastin– and katanin-mediated microtubule disassembly (Lacroix et al., 2010) and also *C. elegans* possesses severing proteins such as a spastin homolog SPAS-1 (Matsushita-Ishiodori et al., 2007) as well as katanin homologs encoded by mei-1 and mei-2 (Lu et al., 2004). Also, elevated tyrosination increases microtubule dynamics (Sanyal et al., 2023), and we have shown that in *ccpp-1* mutants, tubulin tyrosination is enhanced (Fig. 3A+B), likely contributing to the negative effect on motor motility (Fig. 6A+B).

Polyglutamylated tubulin is particularly enriched in the B-tubule of motile cilia and flagella (Lechtreck and Geimer, 2000; Orbach and Howard, 2019), where it plays a key role in regulating dynein activity and ciliary beating (Kubo et al., 2010; Suryavanshi et al., 2010). In *C. elegans,* CCPP and TTLL also express in non-motile cilia such as in ciliated sensory neurons in head, amphids and male-specific tail neurons (Chawla et al., 2016; Kimura et al., 2007; O’Hagan et al., 2011). Interestingly, we identified expression of CCPP enzymes also in ciliated tail-neurons of hermaphrodites as well as in ALM, dorsal cords and sublateral neurons (Fig. 9). More importantly, ccpp-1 promoter reporters significantly colocalize with UNC-104 (Fig. 9A+B) and in Co-IPs, lack of CCPP-1 increases polyglutamylated tubulin binding to UNC-104, while lack of TTLL-5 reduces binding of polyglutamylated tubulin to UNC-104 (Fig. 10A-H). These data from the literature and our own experiments lead to our model (Fig. 11) in which UNC-104 is increasingly trapped on the microtubule surface (due to increased charge-interactions) upon hyperglutamylation, slowing down its motility (Fig. 11B). Impaired axonal transport of synaptic vesicles (Fig. 6H+I) is expected to have physiological consequences for the nematode; and indeed, worm touch responses significantly decreased in *ccpp-1* mutants (Fig. 10 I+J), consistent with findings that tubulin hyperglutamylation is associated with neuropathological defects (Magiera et al., 2018). Thus, *C. elegans* can be used as a model to investigate tubulin glutamylation linked to neurological diseases.

### Impact of hypoglutamylation on UNC-104 motility

Tubulin glutamylation is significantly reduced in *ttll-5(tm3360)* mutants as expected (Fig. 2B+C). TTLL-5 in *C. elegans* is homologous to mammalian TTLL5 (Chawla et al., 2016), which polyglutamylates both α– and β-tubulin, with a preference for α-tubulin. TTLL5 is also capable of initiating and elongating polyglutamylation chains in mammals (van Dijk et al., 2007). Although TTLL-9 in *C. elegans* is homologous to mammalian TTLL9 with polyglutamylase activity (Chawla et al., 2016), it is known that TTLL9 requires interaction with other proteins to acquire enzymatic activity, due to the absence of essential domains (similar to TTLL1 and TTLL2) (Janke et al., 2005). This might explain that simply deleting *ttll-9* alone in *C. elegans* does not cause significant effects on tubulin polyglutamylation (Fig. 2B+C). Indeed, hypoglutamylation enhanced UNC-104 motility in *ttll-5* mutants but was slightly less effective in *ttll-9* mutants (Fig. 8A-C). Also, while *ccpp-1* (inducing hyperglutamylation) leads to visible motor retention in cell bodies (Fig. 7A+B), *ttll-5* (inducing hypoglutamylation) reduces motor presence in neuronal cell bodies (Fig. 8D+E). These results suggest that while hyperglutamylation negatively affects UNC-104-based transport efficiency, hypoglutamylation positively affects UNC-104 motility. Notably, one study shows that hypoglutamylation induced by a TTLL1 deletion in mouse hippocampal neurons results in a twofold increase in mitochondrial motility driven by kinesin-1 (Bodakuntla et al., 2021).

### Crosstalk between different post-translational modifications

Microtubules encounter a wide range of PTMs that regulate their structure and function (Janke and Magiera, 2020). Specifically in dendrites, tubulin PTM largely varies: at the dendritic core, microtubules are generally more acetylated with low levels of tyrosination, while microtubules at the dendritic periphery are more tyrosinated and less acetylated (Tas et al., 2017). Recent findings have demonstrated that vasohibin/SVBP, the enzyme complex responsible for detyrosination, shows markedly reduced activity in the absence of polyglutamylation (Ebberink et al., 2023). Similarly, polyglutamylase TTLL6 exposed minimal activity on unmodified tubulin but 20-fold higher activity on detyrosinated tubulin (Mahalingan et al., 2020). Moreover, in *Ttll1* ^−/−^ *Ttll7* ^−/−^ double mutant mice, high levels of acetylated tubulin are detected (Genova et al., 2023). These findings indicate potential crosstalk between polyglutamylation and other tubulin modifications, including detyrosination and acetylation. Indeed, in *ccpp-1* mutants, tubulin was not only hyperglutamylated (Fig. 1B+C) but also exposed significantly higher tyrosination signals (Fig. 3A+B). Vice versa, deletion of *ttll-5* led not only to hypoglutamylated tubulin (Fig. 2B+C) but also to reduced levels of tyrosination signals (Fig 4A+B). Interestingly, detyrosinated and acetylated tubulin remain unchanged in both mutants (Figs. 3C+D, 4C+D and 5A-D). This finding of unaffected acetylation in *ttll-5* mutants is consistent with a study demonstrating that deletions of *TTLL1* and *TTLL9* in Tetrahymena did not affect tubulin acetylation (Wloga et al., 2008). Because to date no canonical tyrosination enzymes have been characterized in *C. elegans*, we propose that crosstalk between tubulin hyperglutamylation and hypoglutamylation regulates tubulin tyrosination via unknown additional factors in nematodes. This scenario seems more likely than that the glutamylation status directly modulates epitopes recognized by tyrosination antibodies.

### Tubulin tyrosination and its effect on kinesin-3 motility

As mentioned above, kinesin motors “walk” along microtubules by interacting with the C-terminal tails (CTTs) of α– and β-tubulin subunits (Benoit et al., 2024; Lessard et al., 2019). These CTTs are hotspots for PTMs such as tyrosination, detyrosination, acetylation, and polyglutamylation, and these modifications clearly alter the electrostatic and steric environment of the microtubule surface (Janke and Magiera, 2020). Indeed, kinesin motors of the various classes exhibit preferences for specific tubulin modifications. For instance, in mammals, kinesin-3 and kinesin-5 preferentially bind to tyrosinated tubulin enriched on dynamic microtubules near growth cones, distal dendrites and in neuronal cell bodies (Guardia et al., 2016; Kahn et al., 2015). Axonal microtubules, however, are generally enriched in stable forms of tubulin such as acetylated and detyrosinated tubulin, and only the very distal regions near the dynamic growth cones contain more tyrosinated tubulin (Nirschl et al., 2016). Therefore, our finding that reduced tyrosination leads to increased motor motility (Fig. 4A+B and Fig. 8A-C) does not necessarily contrast findings from others that mammalian kinesin-3 prefers tyrosinated tubulin, since our observations are carried out in more proximal regions of axons (as opposed to dynamic growth cones or within neuronal cell bodies). Note that tyrosination does not affect microtubule surface charge (Szczesna et al., 2022); thus, we did not include such surface charge effects in our model in Figure 11.

## ACKNOWLEDGEMENTS

We acknowledge the National Bio-Resource Project (Japan) for kindly providing various strains used in this study. This work was funded by the National Science Council (now NSTC, National Science and Technology Council) under grant number NSC 107-2311-B-007-003-.

## AUTHORS’ CONTRIBUTIONS

OB performed experiments, analyzed data, and wrote the manuscript. MSK and DSP performed experiments and analyzed data. OIW designed experiments, obtained grants, and wrote the manuscript.

## SUPPLEMENTARY FIGURES

**Suppl. Figure S1:**
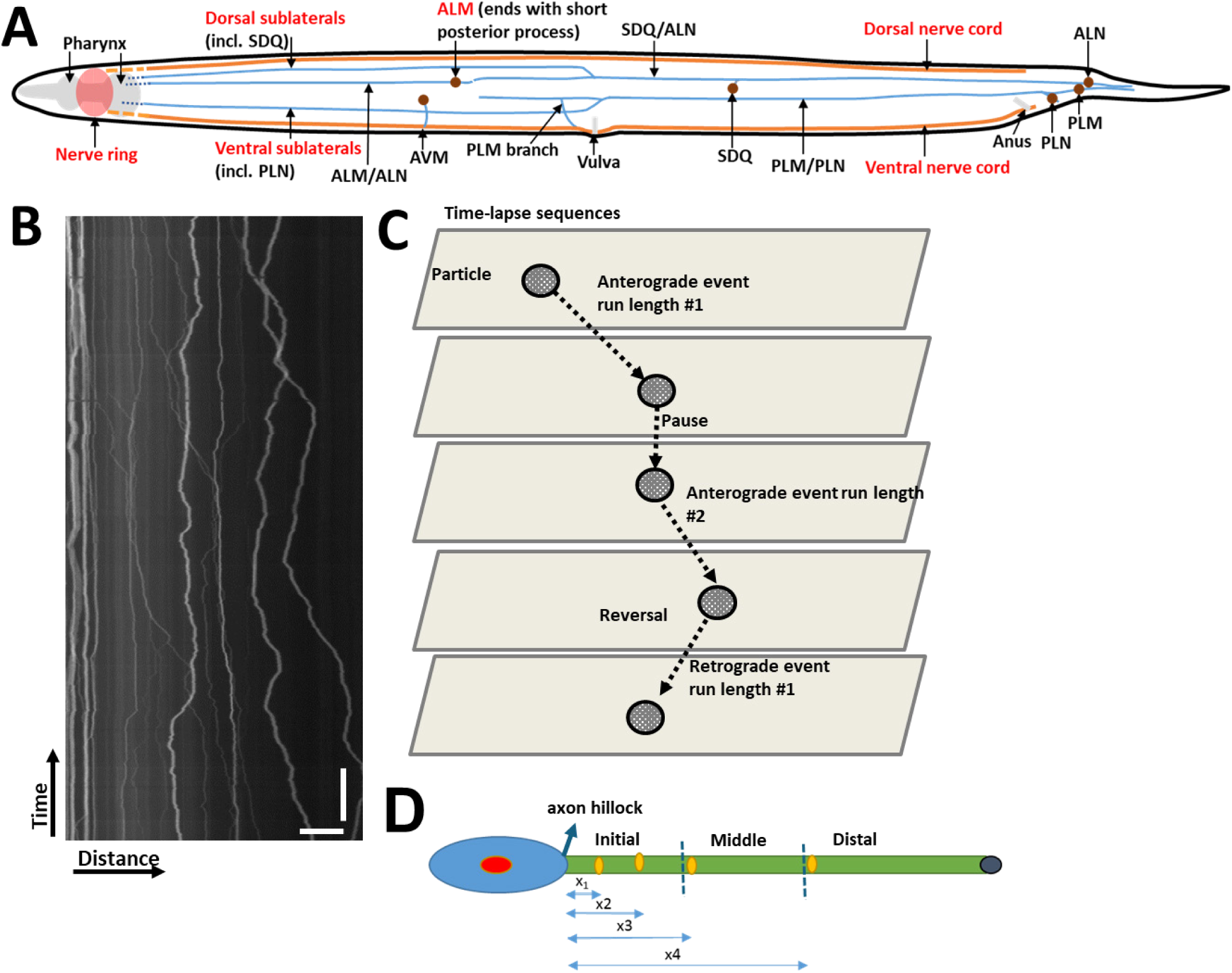
Diagrammatic schemes of mentioned neurons, kymograph analysis as well as cluster analysis methods. (A) Simplified scheme of the nervous system in *C. elegans* (cell bodies in brown color and axons in light blue color). Discussed neurons highlighted in red. (B) Example of a kymograph and (C) diagrammatic scheme of various motility parameters generated from a kymograph. (D) Schematic diagram of a neuron with blue, double-sided arrows indicating particle distances traveled from the axon hillock to farther distal regions. One-sided arrow indicates the relative position of an axon hillock and dotted lines separate the initial, middle and distal regions. Scale bars: vertical = 30 s, horizontal = 10 µm.

**Suppl. Figure S2:**
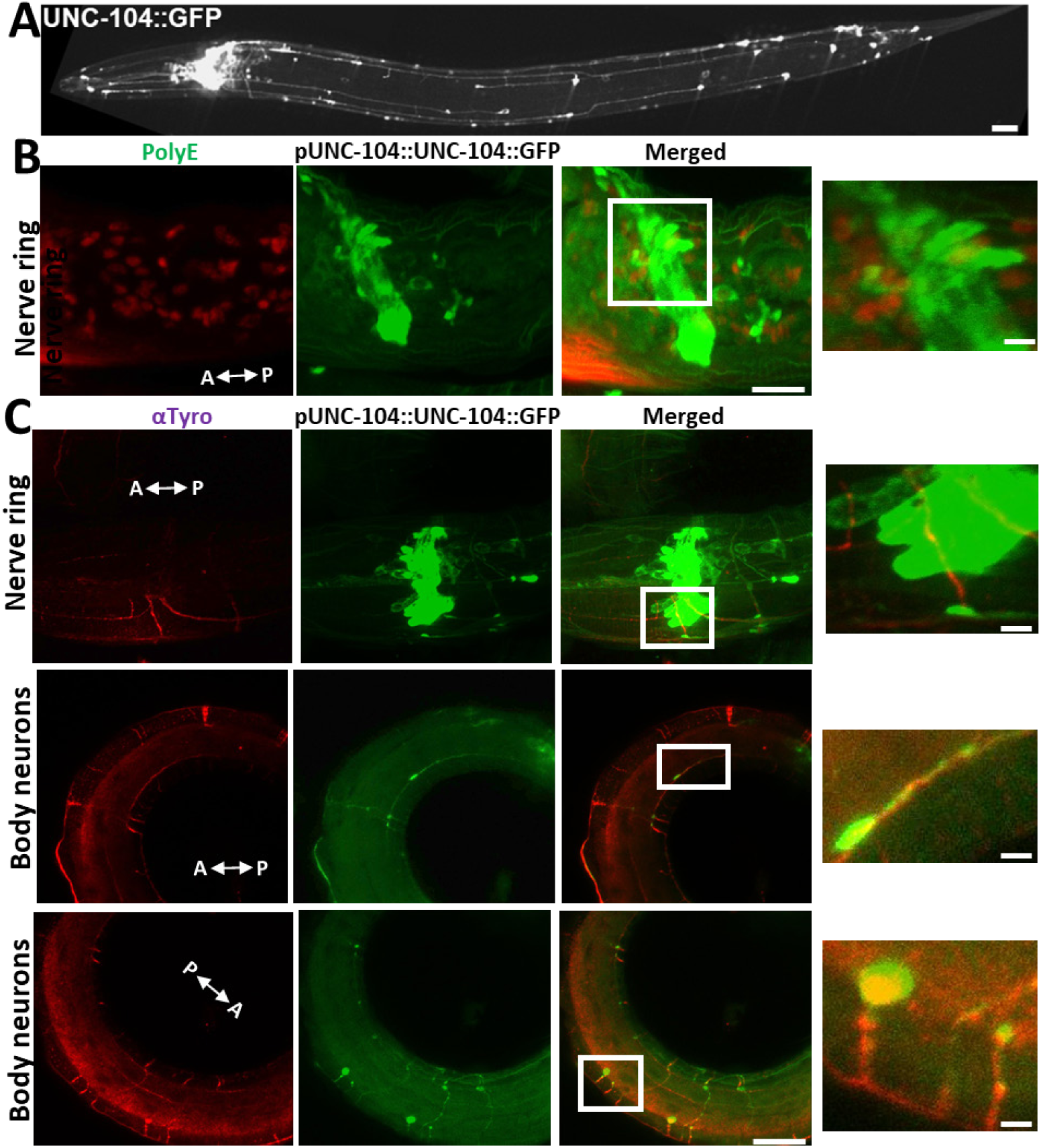
Whole-mount immunostaining reveals polyglutamylated and tyrosinated tubulin colocalizing with UNC-104. (A) Expression of UNC-104::GFP in wild type worms serving as a pan-neuronal marker. (B) Staining of polyglutamylated tubulin in UNC-104::GFP expressing worms. (C) Staining of tyrosinated tubulin in worms expressing UNC-104::GFP. A = Anterior, P = Posterior. White boxes highlight selected regions for the (right-handed) insets. Scale bars: 10 µm; inset: 1 µm

**Suppl. Figure S3:**
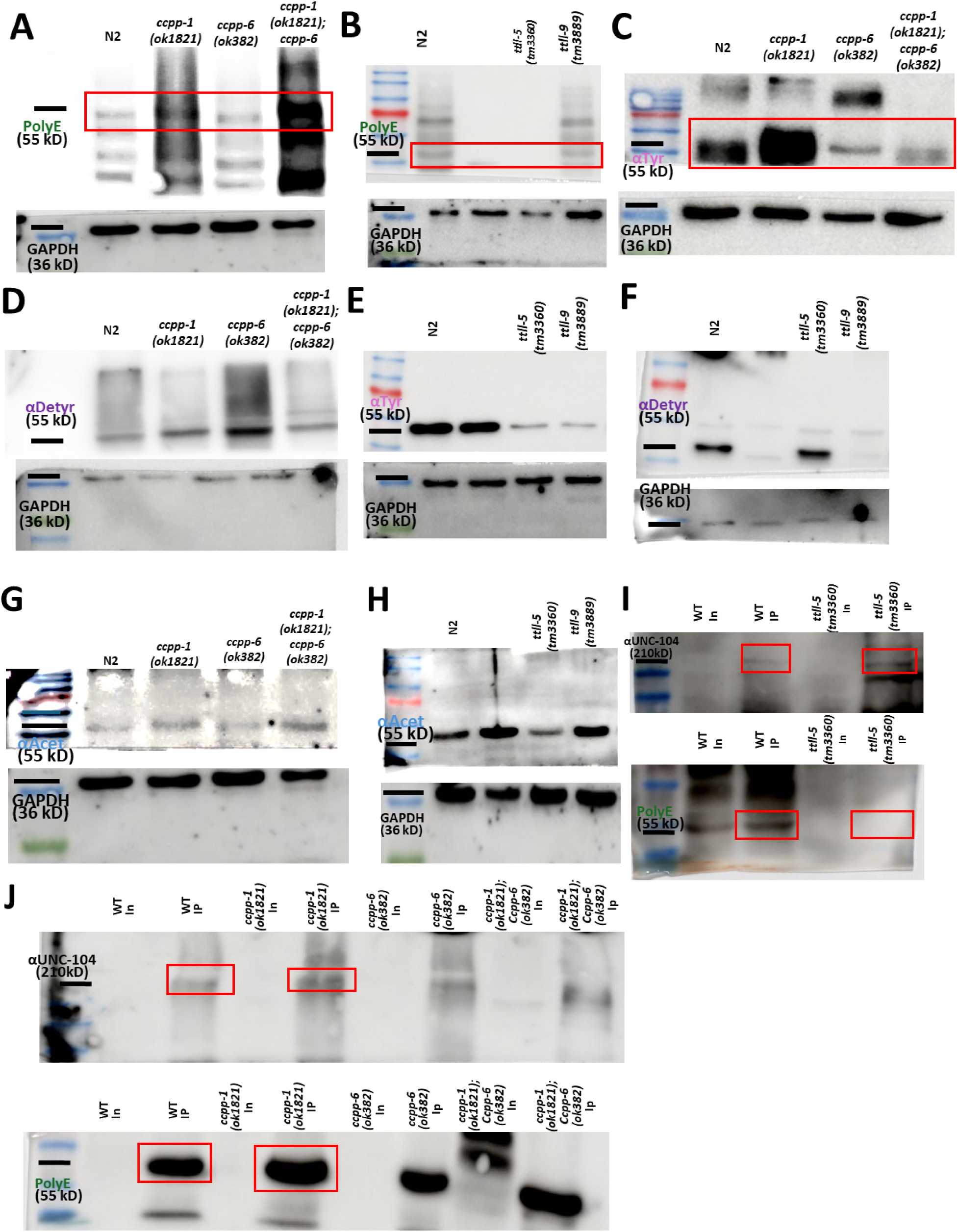
Representative raw images of Western blots and Co-IP (unmodified and boxed). (A) Upper panel: PolyE (55 kD) and lower panel: anti-GAPDH (36 kD) in N2 and deglutamylase mutants (from Fig. 1B). (B) Upper panel: PolyE (55 kD) and Lower panel: anti-GAPDH (36 kD) in N2 and polyglutamylase mutants (from Fig. 2B). (C) Upper panel: αTyrosinated tubulin (55 kD) and Lower panel: anti-GAPDH (36 kD) in N2 and deglutamylase mutants (from Fig. 3A). (D) Upper gel αDetyrosinated tubulin (55 kD) and lower panel αnti-GAPDH (36 kD) in N2 and deglutamylase mutants (from Fig. 3C). (E) Upper gel αTyrosinated tubulin (55 kD) and lower gel αnti-GAPDH (36 kD) in N2 and polgyglutamylase mutants (from Fig. 4A). (F) Upper gel αDetyrosinated tubulin (55 kD) and lower gel αnti-GAPDH (36 kD) in N2 and polgyglutamylase mutants (from Fig. 4C). (G) Upper gel αAcetylated tubulin (55 kD) and lower gel αnti-GAPDH (36 kD) in N2 and deglutamylase mutants (from Fig. 5A). (H) Upper gel αAcetylated tubulin (55 kD) and lower gel αnti-GAPDH (36 kD) in N2 and polyglutamylase mutants (from Fig. 5C). (I) Upper gel αUNC-104 (210 kD) and lower gel PolyE (55 kD) in wild type and *ttll-5(tm3360)* mutants (from Fig. 10A). (J) Upper gel αUNC-104 (210 kD) and lower gel PolyE (55 kD) in wild type and deglutamylase mutants (from Fig. 10E).

## REFERENCES

1. Aillaud, C., C. Bosc, L. Peris, A. Bosson, P. Heemeryck, J. Van Dijk, J. Le Friec, B. Boulan, F. Vossier, L.E. Sanman, S. Syed, N. Amara, Y. Coute, L. Lafanechere, E. Denarier, C. Delphin, L. Pelletier, S. Humbert, M. Bogyo, A. Andrieux, K. Rogowski, and M.J. Moutin. 2017. Vasohibins/SVBP are tubulin carboxypeptidases (TCPs) that regulate neuron differentiation. Science. 358:1448–1453.

2. Bayansan, O., P. Bhan, C.Y. Chang, S.N. Barmaver, C.P. Shen, and O.I. Wagner. 2025. UNC-10/SYD-2 links kinesin-3 to RAB-3-containing vesicles in the absence of the motor’s PH domain. Neurobiol Dis. 204:106766.

3. Benoit, M., L. Rao, A.B. Asenjo, A. Gennerich, and H. Sosa. 2024. Cryo-EM unveils kinesin KIF1A’s processivity mechanism and the impact of its pathogenic variant P305L. Nat Commun. 15:5530.

4. Berezniuk, I., P.J. Lyons, J.J. Sironi, H. Xiao, M. Setou, R.H. Angeletti, K. Ikegami, and L.D. Fricker. 2013. Cytosolic carboxypeptidase 5 removes alpha– and gamma-linked glutamates from tubulin. J Biol Chem. 288:30445–30453.

5. Berezniuk, I., H.T. Vu, P.J. Lyons, J.J. Sironi, H. Xiao, B. Burd, M. Setou, R.H. Angeletti, K. Ikegami, and L.D. Fricker. 2012. Cytosolic carboxypeptidase 1 is involved in processing alpha– and beta-tubulin. J Biol Chem. 287:6503–6517.

6. Bodakuntla, S., X. Yuan, M. Genova, S. Gadadhar, S. Leboucher, M.C. Birling, D. Klein, R. Martini, C. Janke, and M.M. Magiera. 2021. Distinct roles of alpha– and beta-tubulin polyglutamylation in controlling axonal transport and in neurodegeneration. EMBO J. 40:e108498.

7. Bonnet, C., D. Boucher, S. Lazereg, B. Pedrotti, K. Islam, P. Denoulet, and J.C. Larcher. 2001. Differential binding regulation of microtubule-associated proteins MAP1A, MAP1B, and MAP2 by tubulin polyglutamylation. J Biol Chem. 276:12839–12848.

8. Brenner, S. 1974. The genetics of Caenorhabditis elegans. Genetics. 77:71–94.

9. Chalfie, M. 2014. Assaying mechanosensation. WormBook:1–13.

10. Chawla, D.G., R.V. Shah, Z.K. Barth, J.D. Lee, K.E. Badecker, A. Naik, M.M. Brewster, T.P. Salmon, and N. Peel. 2016. Caenorhabditis elegans glutamylating enzymes function redundantly in male mating. Biol Open. 5:1290–1298.

11. Chen, C.W., Y.F. Peng, Y.C. Yen, P. Bhan, M. Muthaiyan Shanmugam, D.R. Klopfenstein, and O.I. Wagner. 2019. Insights on UNC-104-dynein/dynactin interactions and their implications on axonal transport in Caenorhabditis elegans. J Neurosci Res. 97:185–201.

12. Chen, J., and A. Roll-Mecak. 2023. Glutamylation is a negative regulator of microtubule growth. Mol Biol Cell. 34:ar70.

13. Citterio, A., A. Arnoldi, E. Panzeri, L. Merlini, M.G. D’Angelo, O. Musumeci, A. Toscano, A. Bondi, A. Martinuzzi, N. Bresolin, and M.T. Bassi. 2015. Variants in KIF1A gene in dominant and sporadic forms of hereditary spastic paraparesis. Journal of neurology. 262:2684–2690.

14. Dominguez, J., B.P. Shah, and N. Peel. 2022. A ccpp-6 deletion mutation does not impair gross cilia integrity in C. elegans. MicroPubl Biol. 2022.

15. Dunn, S., E.E. Morrison, T.B. Liverpool, C. Molina-Paris, R.A. Cross, M.C. Alonso, and M. Peckham. 2008. Differential trafficking of Kif5c on tyrosinated and detyrosinated microtubules in live cells. J Cell Sci. 121:1085–1095.

16. Ebberink, E., S. Fernandes, G. Hatzopoulos, N. Agashe, P.H. Chang, N. Guidotti, T.M. Reichart, L. Reymond, M.C. Velluz, F. Schneider, C. Pourroy, C. Janke, P. Gonczy, B. Fierz, and C. Aumeier. 2023. Tubulin engineering by semi-synthesis reveals that polyglutamylation directs detyrosination. Nat Chem. 15:1179–1187.

17. Eves-van den Akker, S., C.J. Lilley, J.R. Ault, A.E. Ashcroft, J.T. Jones, and P.E. Urwin. 2014. The feeding tube of cyst nematodes: characterisation of protein exclusion. PLoS One. 9:e87289.

18. Fernandez-Gonzalez, A., A.R. La Spada, J. Treadaway, J.C. Higdon, B.S. Harris, R.L. Sidman, J.I. Morgan, and J. Zuo. 2002. Purkinje cell degeneration (pcd) phenotypes caused by mutations in the axotomy-induced gene, Nna1. Science. 295:1904–1906.

19. Genova, M., L. Grycova, V. Puttrich, M.M. Magiera, Z. Lansky, C. Janke, and M. Braun. 2023. Tubulin polyglutamylation differentially regulates microtubule-interacting proteins. EMBO J. 42:e112101.

20. Ghosh-Roy, A., A. Goncharov, Y. Jin, and A.D. Chisholm. 2012. Kinesin-13 and tubulin posttranslational modifications regulate microtubule growth in axon regeneration. Dev Cell. 23:716–728.

21. Guardia, C.M., G.G. Farias, R. Jia, J. Pu, and J.S. Bonifacino. 2016. BORC Functions Upstream of Kinesins 1 and 3 to Coordinate Regional Movement of Lysosomes along Different Microtubule Tracks. Cell Rep. 17:1950–1961.

22. Hall, D.H., and E.M. Hedgecock. 1991. Kinesin-related gene unc-104 is required for axonal transport of synaptic vesicles in C. elegans. Cell. 65:837–847.

23. Hancock, W.O. 2014. Bidirectional cargo transport: moving beyond tug of war. Nat Rev Mol Cell Biol. 15:615–628.

24. Hotta, T., A. Plemmons, M. Gebbie, T.A. Ziehm, T.L. Blasius, C. Johnson, K.J. Verhey, J.N. Pearring, and R. Ohi. 2023. Mechanistic Analysis of CCP1 in Generating DeltaC2 alpha-Tubulin in Mammalian Cells and Photoreceptor Neurons. Biomolecules. 13.

25. Ikegami, K., R.L. Heier, M. Taruishi, H. Takagi, M. Mukai, S. Shimma, S. Taira, K. Hatanaka, N. Morone, I. Yao, P.K. Campbell, S. Yuasa, C. Janke, G.R. Macgregor, and M. Setou. 2007. Loss of alpha-tubulin polyglutamylation in ROSA22 mice is associated with abnormal targeting of KIF1A and modulated synaptic function. Proc Natl Acad Sci U S A. 104:3213–3218.

26. Janke, C., and M.M. Magiera. 2020. The tubulin code and its role in controlling microtubule properties and functions. Nat Rev Mol Cell Biol. 21:307–326.

27. Janke, C., K. Rogowski, and J. van Dijk. 2008. Polyglutamylation: a fine-regulator of protein function? ‘Protein Modifications: beyond the usual suspects’ review series. EMBO Rep. 9:636–641.

28. Janke, C., K. Rogowski, D. Wloga, C. Regnard, A.V. Kajava, J.M. Strub, N. Temurak, J. van Dijk, D. Boucher, A. van Dorsselaer, S. Suryavanshi, J. Gaertig, and B. Edde. 2005. Tubulin polyglutamylase enzymes are members of the TTL domain protein family. Science. 308:1758–1762.

29. Kahn, O.I., V. Sharma, C. Gonzalez-Billault, and P.W. Baas. 2015. Effects of kinesin-5 inhibition on dendritic architecture and microtubule organization. Mol Biol Cell. 26:66–77.

30. Kalinina, E., R. Biswas, I. Berezniuk, A. Hermoso, F.X. Aviles, and L.D. Fricker. 2007. A novel subfamily of mouse cytosolic carboxypeptidases. FASEB J. 21:836–850.

31. Kimura, S., T. Noda, and T. Yoshimori. 2007. Dissection of the autophagosome maturation process by a novel reporter protein, tandem fluorescent-tagged LC3. Autophagy. 3:452–460.

32. Kimura, Y., N. Kurabe, K. Ikegami, K. Tsutsumi, Y. Konishi, O.I. Kaplan, H. Kunitomo, Y. Iino, O.E. Blacque, and M. Setou. 2010. Identification of tubulin deglutamylase among Caenorhabditis elegans and mammalian cytosolic carboxypeptidases (CCPs). J Biol Chem. 285:22936–22941.

33. Klimas, A.S., J. Dominguez, B.P. Shah, Z.Y. Lee, and N. Peel. 2023. The C. elegans deglutamylase CCPP-6 does not operate redundantly with CCPP-1 in gross cilia integrity. MicroPubl Biol. 2023.

34. Kravec, M., O. Šedo, J. Nedvědová, M. Micka, M. Šulcová, N. Zezula, K. Gömöryová, D. Potěšil, R. Sri Ganji, S. Bologna, I. Červenka, Z. Zdráhal, J. Harnoš, K. Tripsianes, C. Janke, C. Bařinka, and V. Bryja. 2024. Carboxy-terminal polyglutamylation regulates signaling and phase separation of the Dishevelled protein. The EMBO Journal. 43:5635–5666.

35. Kubo, T., H.A. Yanagisawa, T. Yagi, M. Hirono, and R. Kamiya. 2010. Tubulin polyglutamylation regulates axonemal motility by modulating activities of inner-arm dyneins. Curr Biol. 20:441–445.

36. Lacroix, B., J. van Dijk, N.D. Gold, J. Guizetti, G. Aldrian-Herrada, K. Rogowski, D.W. Gerlich, and C. Janke. 2010. Tubulin polyglutamylation stimulates spastin-mediated microtubule severing. J Cell Biol. 189:945–954.

37. Lechtreck, K.F., and S. Geimer. 2000. Distribution of polyglutamylated tubulin in the flagellar apparatus of green flagellates. Cell Motil Cytoskeleton. 47:219–235.

38. Lessard, D.V., O.J. Zinder, T. Hotta, K.J. Verhey, R. Ohi, and C.L. Berger. 2019. Polyglutamylation of tubulin’s Cterminal tail controls pausing and motility of kinesin-3 family member KIF1A. J Biol Chem. 294:6353–6363.

39. Li, L., and X.J. Yang. 2015. Tubulin acetylation: responsible enzymes, biological functions and human diseases. Cell Mol Life Sci. 72:4237–4255.

40. Lu, C., M. Srayko, and P.E. Mains. 2004. The Caenorhabditis elegans microtubule-severing complex MEI-1/MEI-2 katanin interacts differently with two superficially redundant beta-tubulin isotypes. Mol Biol Cell. 15:142–150.

41. Lu, Y.M., and C. Zheng. 2022. The Expression and Function of Tubulin Isotypes in Caenorhabditis elegans. Front Cell Dev Biol. 10:860065.

42. Magiera, M.M., S. Bodakuntla, J. Ziak, S. Lacomme, P. Marques Sousa, S. Leboucher, T.J. Hausrat, C. Bosc, A. Andrieux, M. Kneussel, M. Landry, A. Calas, M. Balastik, and C. Janke. 2018. Excessive tubulin polyglutamylation causes neurodegeneration and perturbs neuronal transport. EMBO J. 37.

43. Mahalingan, K.K., E. Keith Keenan, M. Strickland, Y. Li, Y. Liu, H.L. Ball, M.E. Tanner, N. Tjandra, and A. Roll-Mecak. 2020. Structural basis for polyglutamate chain initiation and elongation by TTLL family enzymes. Nat Struct Mol Biol. 27:802–813.

44. Matsushita-Ishiodori, Y., K. Yamanaka, and T. Ogura. 2007. The C. elegans homologue of the spastic paraplegia protein, spastin, disassembles microtubules. Biochem Biophys Res Commun. 359:157–162.

45. Millecamps, S., and J.P. Julien. 2013. Axonal transport deficits and neurodegenerative diseases. Nat Rev Neurosci. 14:161–176.

46. Morikawa, M., N.U. Jerath, T. Ogawa, M. Morikawa, Y. Tanaka, M.E. Shy, S. Zuchner, and N. Hirokawa. 2022. A neuropathy-associated kinesin KIF1A mutation hyper-stabilizes the motor-neck interaction during the ATPase cycle. Embo j. 41:e108899.

47. Moutin, M.J., C. Bosc, L. Peris, and A. Andrieux. 2021. Tubulin post-translational modifications control neuronal development and functions. Dev Neurobiol. 81:253–272.

48. Muniesh, M.S., S.N. Barmaver, H.Y. Huang, O. Bayansan, and O.I. Wagner. 2020. PTP-3 phosphatase promotes intramolecular folding of SYD-2 to inactivate kinesin-3 UNC-104 in neurons. Mol Biol Cell. 31:2932–2947.

49. Neumann, B., and M.A. Hilliard. 2014. Loss of MEC-17 leads to microtubule instability and axonal degeneration. Cell Rep. 6:93–103.

50. Nirschl, J.J., M.M. Magiera, J.E. Lazarus, C. Janke, and E.L. Holzbaur. 2016. alpha-Tubulin Tyrosination and CLIP-170 Phosphorylation Regulate the Initiation of Dynein-Driven Transport in Neurons. Cell Rep. 14:2637–2652.

51. O’Hagan, R., B.P. Piasecki, M. Silva, P. Phirke, K.C. Nguyen, D.H. Hall, P. Swoboda, and M.M. Barr. 2011. The tubulin deglutamylase CCPP-1 regulates the function and stability of sensory cilia in C. elegans. Curr Biol. 21:1685–1694.

52. Orbach, R., and J. Howard. 2019. The dynamic and structural properties of axonemal tubulins support the high length stability of cilia. Nat Commun. 10:1838.

53. Pagnamenta, A.T., P. Heemeryck, H.C. Martin, C. Bosc, L. Peris, I. Uszynski, S. Gory-Faure, S. Couly, C. Deshpande, A. Siddiqui, A.A. Elmonairy, W.G.S. Consortium, C. Genomics England Research, S. Jayawant, S. Murthy, I. Walker, L. Loong, P. Bauer, F. Vossier, E. Denarier, T. Maurice, E.L. Barbier, J.C. Deloulme, J.C. Taylor, E.M. Blair, A. Andrieux, and M.J. Moutin. 2019. Defective tubulin detyrosination causes structural brain abnormalities with cognitive deficiency in humans and mice. Hum Mol Genet. 28:3391–3405.

54. Rogowski, K., J. van Dijk, M.M. Magiera, C. Bosc, J.C. Deloulme, A. Bosson, L. Peris, N.D. Gold, B. Lacroix, M. Bosch Grau, N. Bec, C. Larroque, S. Desagher, M. Holzer, A. Andrieux, M.J. Moutin, and C. Janke. 2010. A family of protein-deglutamylating enzymes associated with neurodegeneration. Cell. 143:564–578.

55. Sanyal, C., N. Pietsch, S. Ramirez Rios, L. Peris, L. Carrier, and M.J. Moutin. 2023. The detyrosination/retyrosination cycle of tubulin and its role and dysfunction in neurons and cardiomyocytes. Semin Cell Dev Biol. 137:46–62.

56. Schneider, C.A., W.S. Rasband, and K.W. Eliceiri. 2012. NIH Image to ImageJ: 25 years of image analysis. Nat Methods. 9:671–675.

57. Soppina, V., and K.J. Verhey. 2014. The family-specific K-loop influences the microtubule on-rate but not the superprocessivity of kinesin-3 motors. Mol Biol Cell. 25:2161–2170.

58. Suryavanshi, S., B. Edde, L.A. Fox, S. Guerrero, R. Hard, T. Hennessey, A. Kabi, D. Malison, D. Pennock, W.S. Sale, D. Wloga, and J. Gaertig. 2010. Tubulin glutamylation regulates ciliary motility by altering inner dynein arm activity. Curr Biol. 20:435–440.

59. Szczesna, E., E.A. Zehr, S.W. Cummings, A. Szyk, K.K. Mahalingan, Y. Li, and A. Roll-Mecak. 2022. Combinatorial and antagonistic effects of tubulin glutamylation and glycylation on katanin microtubule severing. Dev Cell. 57:2497–2513 e2496.

60. Tas, R.P., A. Chazeau, B.M.C. Cloin, M.L.A. Lambers, C.C. Hoogenraad, and L.C. Kapitein. 2017. Differentiation between Oppositely Oriented Microtubules Controls Polarized Neuronal Transport. Neuron. 96:1264–1271 e1265.

61. Tien, N.W., G.H. Wu, C.C. Hsu, C.Y. Chang, and O.I. Wagner. 2011. Tau/PTL-1 associates with kinesin-3 KIF1A/UNC-104 and affects the motor’s motility characteristics in C. elegans neurons. Neurobiol Dis. 43:495–506.

62. Valenstein, M.L., and A. Roll-Mecak. 2016. Graded Control of Microtubule Severing by Tubulin Glutamylation. Cell. 164:911–921.

63. van Dijk, J., K. Rogowski, J. Miro, B. Lacroix, B. Edde, and C. Janke. 2007. A targeted multienzyme mechanism for selective microtubule polyglutamylation. Mol Cell. 26:437–448.

64. Wagner, O.I., A. Esposito, B. Kohler, C.W. Chen, C.P. Shen, G.H. Wu, E. Butkevich, S. Mandalapu, D. Wenzel, F.S. Wouters, and D.R. Klopfenstein. 2009. Synaptic scaffolding protein SYD-2 clusters and activates kinesin-3 UNC-104 in C. elegans. Proc Natl Acad Sci U S A. 106:19605–19610.

65. Wang, X., K. Song, Y. Li, L. Tang, and X. Deng. 2019. Single-Molecule Imaging and Computational Microscopy Approaches Clarify the Mechanism of the Dimerization and Membrane Interactions of Green Fluorescent Protein. Int J Mol Sci. 20.

66. Ward, S., D.J. Burke, J.E. Sulston, A.R. Coulson, D.G. Albertson, D. Ammons, M. Klass, and E. Hogan. 1988. Genomic organization of major sperm protein genes and pseudogenes in the nematode Caenorhabditis elegans. J Mol Biol. 199:1–13.

67. Wloga, D., K. Rogowski, N. Sharma, J. Van Dijk, C. Janke, B. Edde, M.H. Bre, N. Levilliers, V. Redeker, J. Duan, M.A. Gorovsky, M. Jerka-Dziadosz, and J. Gaertig. 2008. Glutamylation on alpha-tubulin is not essential but affects the assembly and functions of a subset of microtubules in Tetrahymena thermophila. Eukaryot Cell. 7:1362–1372.

68. Wu, G.H., M. Muthaiyan Shanmugam, P. Bhan, Y.H. Huang, and O.I. Wagner. 2016. Identification and Characterization of LIN-2(CASK) as a Regulator of Kinesin-3 UNC-104(KIF1A) Motility and Clustering in Neurons. Traffic. 17:891–907.

69. Wu, H.Y., Y. Rong, K. Correia, J. Min, and J.I. Morgan. 2015. Comparison of the enzymatic and functional properties of three cytosolic carboxypeptidase family members. J Biol Chem. 290:1222–1232.

70. Yonekawa, Y., A. Harada, Y. Okada, T. Funakoshi, Y. Kanai, Y. Takei, S. Terada, T. Noda, and N. Hirokawa. 1998. Defect in synaptic vesicle precursor transport and neuronal cell death in KIF1A motor protein-deficient mice. J Cell Biol. 141:431–441.

71. Zaniewski, T.M., and W.O. Hancock. 2023. Positive charge in the K-loop of the kinesin-3 motor KIF1A regulates superprocessivity by enhancing microtubule affinity in the one-head-bound state. J Biol Chem. 299:102818.

